# Citrate Compartmentalization Controls Calcium-Dependent Cytokine Production in Effector T Cells

**DOI:** 10.64898/2026.06.11.731694

**Authors:** Andrea L. Cote, Claire L. McIntyre, Joshua A. Acklin, Lillian R. Delacruz, Yunping Qiu, Irwin J. Kurland, Alison E. Ringel

**Affiliations:** Ragon Institute of Mass General Brigham, MIT, and Harvard, Cambridge, MA 02139, USA; MIT Biology Department, Cambridge, MA 02139, USA; Albert Einstein College of Medicine, Fleischer Institute for Diabetes and Metabolism, Department of Medicine, Division of Endocrinology, Bronx, NY USA 10461; Stable Isotope and Metabolomics Core Facility, New York Regional Diabetes Research Center at the Albert Einstein College of Medicine, Bronx, NY, USA 10461; Koch Institute for Integrative Cancer Research, Cambridge, MA 02139, USA

## Abstract

Cytokine production is a core function of effector T cells, yet the mechanisms that regulate cytokine output during an immune response remain incompletely understood. Here, we identify citrate compartmentalization as a cellular mechanism by which CD8^+^ T cells couple cytokine production to glucose availability. Under glucose-replete conditions, citrate transport from the mitochondria to the cytosol by the citrate carrier SLC25A1 suppresses calcium-dependent transcription factor activity in effector T cells. Either reducing glucose availability or blocking the exchange of citrate across the mitochondrial membrane raises free cytosolic calcium, thereby driving nuclear localization of Nuclear Factor of Activated T cells (NFAT)-family transcription factors and sustaining cytokine production. As a calcium-chelating metabolite, we show that citrate buffers free cytosolic calcium, thereby linking calcium-dependent signaling to mitochondrial fuel oxidation. We also identify signatures of this regulatory mechanism across hundreds of human cancer cell lines, where there are negative associations between citrate-derived metabolites and calcium-dependent transcriptional programs, and within the spatial organization of human tumors. These findings identify cytosolic citrate as a broadly conserved metabolic rheostat coupling glucose availability to calcium signaling. By adding calcium signaling to the known functions regulated by SLC25A1, our work reveals a mechanism by which mitochondria adaptively tune cytokine expression and other calcium-dependent programs in response to local metabolic conditions, such as nutrients that are available within a tissue or tumor.

## Introduction

During activation, CD8^+^ T cells transition from a naïve to an effector state, acquiring the ability to produce inflammatory cytokines and perform targeted killing of abnormal cells. This process is coordinated by surface receptors that engage signaling networks converging on a core set of transcription factors, which collectively remodel the transcriptome and proteome. In addition to endowing T cells with functions necessary for host protection, these signals also increase the uptake of extracellular fuels [1–4], drive mitochondrial biogenesis [5,6], and broadly reprogram metabolism [7,8]. Metabolic reprogramming produces biomolecules that sustain proliferation and bioenergetic demands associated with activation, while also producing metabolic signals that reinforce effector differentiation [7,9,10]. The recognition that cellular metabolism and canonical activation signals interact bidirectionally in T cells has revealed that many effector functions are also under metabolic control [7,11].

The observation that activation signals induce glucose metabolism was among the earliest metabolic findings in T lymphocytes [12–15] and is now understood to coordinate an array of cellular programs required for effector responses. Co-stimulation induces glucose uptake [2,3], which rises rapidly as T cells activate [1,2,16] and remains at high levels throughout the effector phase. While glycolysis generates ATP and biomass needed to support rapid proliferation, intermediates derived from glycolysis also serve as signals that commit activating T cells to effector fates [17–19]. During early activation, glycolysis provides ATP that sustains PI3K-Akt signaling [20,21]. Early glycolytic reprogramming also generates phosphoenolpyruvate, which reinforces calcium-dependent transcription by NFAT (Nuclear Factor of Activated T cells) by slowing the removal of calcium from the cytosol by SERCA channels [22,23]. Glycolytic metabolism is essential for T cell activation, as complete glucose withdrawal or severe restriction abrogates proliferation and cytokine production [24–29]. However, studies where glucose has been manipulated in less restrictive ways have revealed more modular and nuanced dependencies, where some functions require aerobic glycolysis, but others require mitochondrial glucose oxidation [30]. Rebalancing how glucose is used during activation can confer functional benefits to T cells as well. Augmenting mitochondrial glucose oxidation, which generates citrate as the first step in the Tricarboxylic Acid (TCA) cycle [31], as well as transiently reducing glucose availability during activation, both enhance anti-tumor efficacy in adoptive transfer models. Thus, the availability of glucose and how it is metabolized regulate effector programs during T cell activation, but the mechanisms are still incompletely understood.

While having sufficient access to extracellular fuels and nutrients is clearly necessary for T cell activation and survival, far less is known about how natural variation in nutrient availability instructs the acquisition of specialized effector functions. Here, we have identified mitochondrial glucose metabolism as a rheostat that controls calcium-dependent signaling processes through citrate export to the cytosol. In glucose-replete conditions, effector T cells transfer citrate from the mitochondria to the cytosol, which buffers free calcium and excludes NFAT from the nucleus, a calcium-dependent transcription factor that controls cytokine expression. Moderate glucose restriction favors its complete oxidation by the TCA cycle, leading to retention of citrate in the mitochondria that promotes expression of NFAT target genes. More broadly, we identify significant inverse associations between calcium-dependent transcriptional programs and citrate-related metabolites across the entire Cancer Cell Line Encyclopedia (CCLE) [32] and within human patient tumor samples. Thus, our work reveals citrate compartmentalization as a regulatory mechanism linking mitochondrial metabolism to calcium-dependent cellular functions, such as cytokine production in effector T cells.

## Results

### Moderate glucose limitation enhances cytokine production during T cell activation

Given that withdrawing glucose has been previously shown to impair a wide range of T cell functions important for host protection [2,24–26,29,33–35], we sought to understand how effector programs instead respond to more moderate glucose limitation. We chose to compare T cells stimulated under glucose concentrations matching conventional RPMI medium (10 mM glucose, referred to as “high”) with those cultured in limited glucose (1.5 mM, referred to as “low”) across three days of polyclonal activation on anti-CD3 and anti-CD28 antibodies. These concentrations were selected to bracket a range of glucose levels with physiological correlates *in vivo*. Healthy adults maintain fasting plasma glucose concentrations of ∼5.5 mM [36,37], whereas blood glucose in individuals with uncontrolled type 2 diabetes can exceed 10 mM [38]. At the lower end, glucose in the interstitial fluid of skeletal muscle and adipose tissue has been reported to be up to 40% lower than in plasma [39–41], and intratumoral glucose concentrations can fall below 1 mM in some tumors due to high local metabolic demand [42]. We evaluated the capacity to produce cytokines following phorbol 12-myristate 13-acetate (PMA)/ ionomycin stimulation across a three-day activation time course by flow cytometry (**Figure 1A**). This revealed that moderate glucose limitation sustained high levels of Interleukin-2 (IL-2) production across the full 72 hours of activation (**Figures 1B-D**), even though IL-2 production in high glucose declines substantially over this time frame [43,44]. A similar pattern was observed for other inflammatory cytokines, including Interferon Gamma (IFN-γ) (**Figures 1E-G**) and Tumor Necrosis Factor Alpha (TNF-α) (**Figures 1H-J**). A larger proportion of T cells were polyfunctional when activated in low glucose, expressing all three cytokines simultaneously (**Figure 1K**). However, this gain in cytokine functionality was not coupled to other effector programs. Low glucose still supported proliferation, but there were fewer cell divisions between two and three days of stimulation compared to the high glucose condition (**Figures S1A-D**). T cells activated under mild glucose restriction expressed slightly less Granzyme B (**Figure 1L**) but displayed equivalent markers of activation, including PD-1 (**Figure 1M**), CD69 (**Figures S1E-F**), and CD25 (**Figures S1G-H**). There were subtle shifts in differentiation markers as well, with a slight decrease in central memory T cells (**Figure S1I**) and a concomitant increase in cells bearing effector memory markers (**Figure S1J**). Overall, we find that T cells activated in low glucose have a greater capacity to produce cytokines, with the largest effect on IL-2.

**Figure 1:**
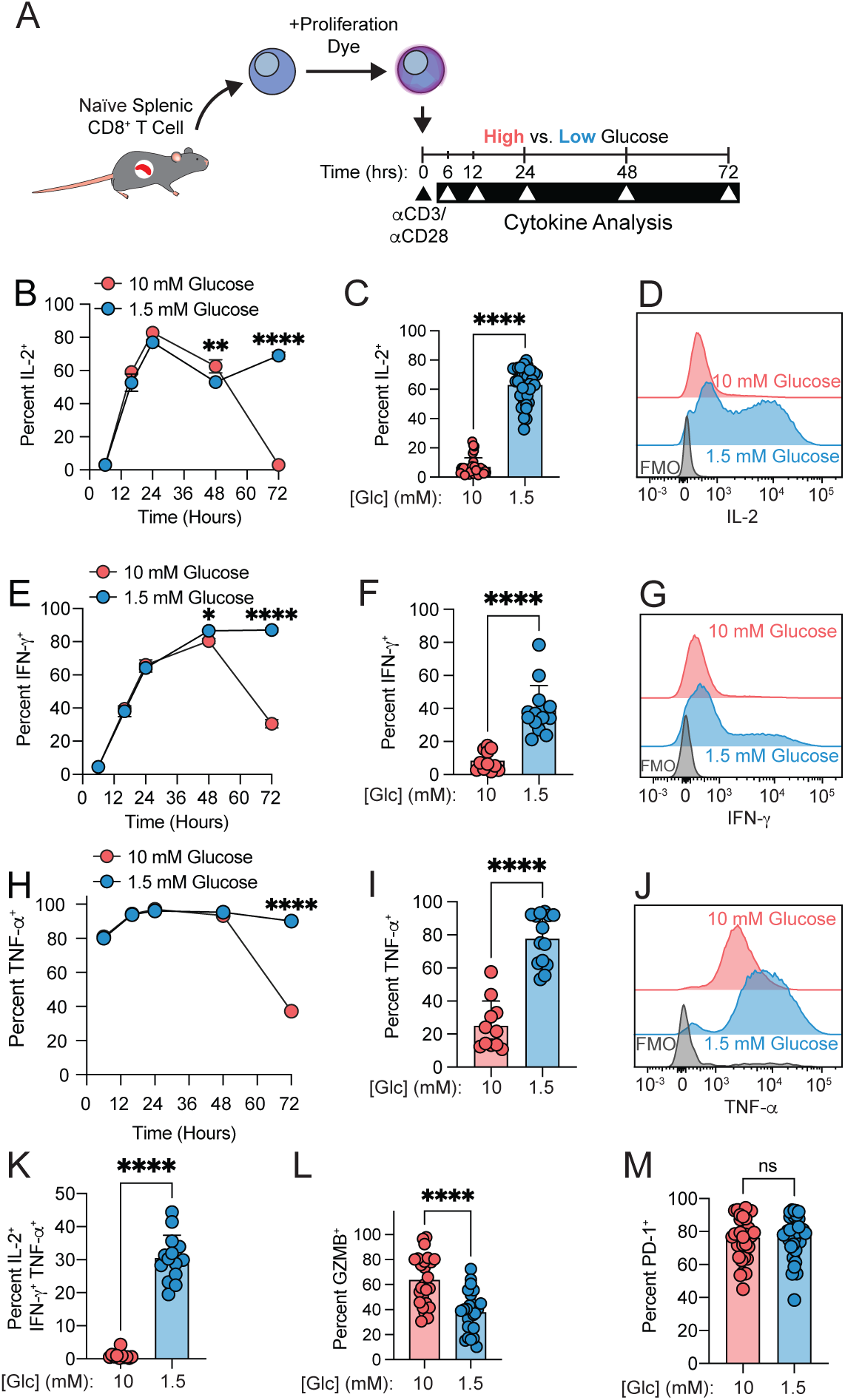
Moderate glucose limitation maintains high cytokine production during T cell activation. (**A**) Experimental schematic for Panels B-M. (**B-D**) Percent IL-2^+^ of live CD8^+^ T cells at multiple time points of polyclonal stimulation by plate-bound α-CD3/ α-CD28 in high (10 mM) versus low (1.5 mM) glucose (**B**), or after 72 hours (**C**). Representative histograms after 72 hours of activation (**D**). (**E-G**) Percent IFN-γ^+^ of live CD8^+^ T cells at multiple time points of polyclonal stimulation by plate-bound α-CD3/ α-CD28 in high (10 mM) versus low (1.5 mM) glucose (**E**), or after 72 hours (**F**). Representative histograms after 72 hours of activation (**G**). (**H-J**) Percent TNF-α^+^ of live CD8^+^ T cells at multiple time points of polyclonal stimulation by plate-bound α-CD3/ α-CD28 in high (10 mM) versus low (1.5 mM) glucose (**H**), or after 72 hours (**I**). Representative histograms after 72 hours of activation (**J**). (**K-M**) Barplots depicting the percent polyfunctional (IL-2^+^ IFN-γ^+^ TNF-α^+^) of live CD8^+^ T cells (**K**), GZMB^+^ of live CD8^+^ T cells (**L**), or PD-1^+^ of live CD8^+^ T cells (**M**) after 72 hours of activation in high (10 mM) versus low (1.5 mM) glucose. *Abbreviations*: Glc = glucose; FMO = Fluorescence Minus One (control condition); IFN-γ = Interferon Gamma; TNF-α = Tumor Necrosis Factor alpha; GZMB = Granzyme B; PD-1 = Programmed Cell Death Protein 1 *Statistics*: All data points are distinct biological replicates. Data are presented as mean ± standard deviation (B, C, E, F, H, I, K-M). Panels B, E, and H show data representative of two independent experiments where n=3 biological replicates are shown from a representative experiment. Panel C contains data aggregated from 10 independent experiments with n=34 biological replicates. Panels F, I, and K-M contains data aggregated from 10 independent experiments with n=22 biological replicates. Statistical significance assessed by Student’s t-test (B, C, E, F, H, I, K-M). P-values: not significant (ns) for p>0.05, ∗ for p≤0.05, ∗∗ for p≤0.01, ∗∗∗ for p≤0.001, and ∗∗∗∗ for p≤0.0001.

We next asked whether the enhanced cytokine production observed under glucose limitation was conserved across different T cell activation conditions. First, we evaluated a model of antigen-specific activation by stimulating OT-I CD8^+^ T cells, which express a transgenic T Cell Receptor (TCR) that recognizes a peptide derived from hen egg white ovalbumin [45], with cognate peptide (**Figure 2A**). After three days in culture, we observed higher IL-2 (**Figure 2B**), IFN-γ (**Figure 2C**), and TNF-α (**Figure 2D**) expression, as well as more polyfunctional T cells (**Figure 2E**) under glucose limitation. Next, we determined how the duration of the activation signal impacted cytokine expression. CD8^+^ T cells were stimulated with plate-bound anti-CD3/anti-CD28 for 24 hours and then expanded in IL-2 cytokine for another three days. Glucose restriction still enhanced the expression of all three cytokines with transient TCR stimulation and expansion (**Figures 2F-J**). In both cases, glucose limitation reduced proliferation (**Figures S2A-B**). Thus, glucose limitation consistently raises the capacity to produce inflammatory cytokines regardless of the activation cue. As these studies were conducted in primary mouse T cells, we next asked whether this effect was conserved in human T cells. Likewise, reducing glucose availability significantly raised polyfunctional cytokine production in human CD8^+^ T cells (**Figure 2K-O**).

**Figure 2:**
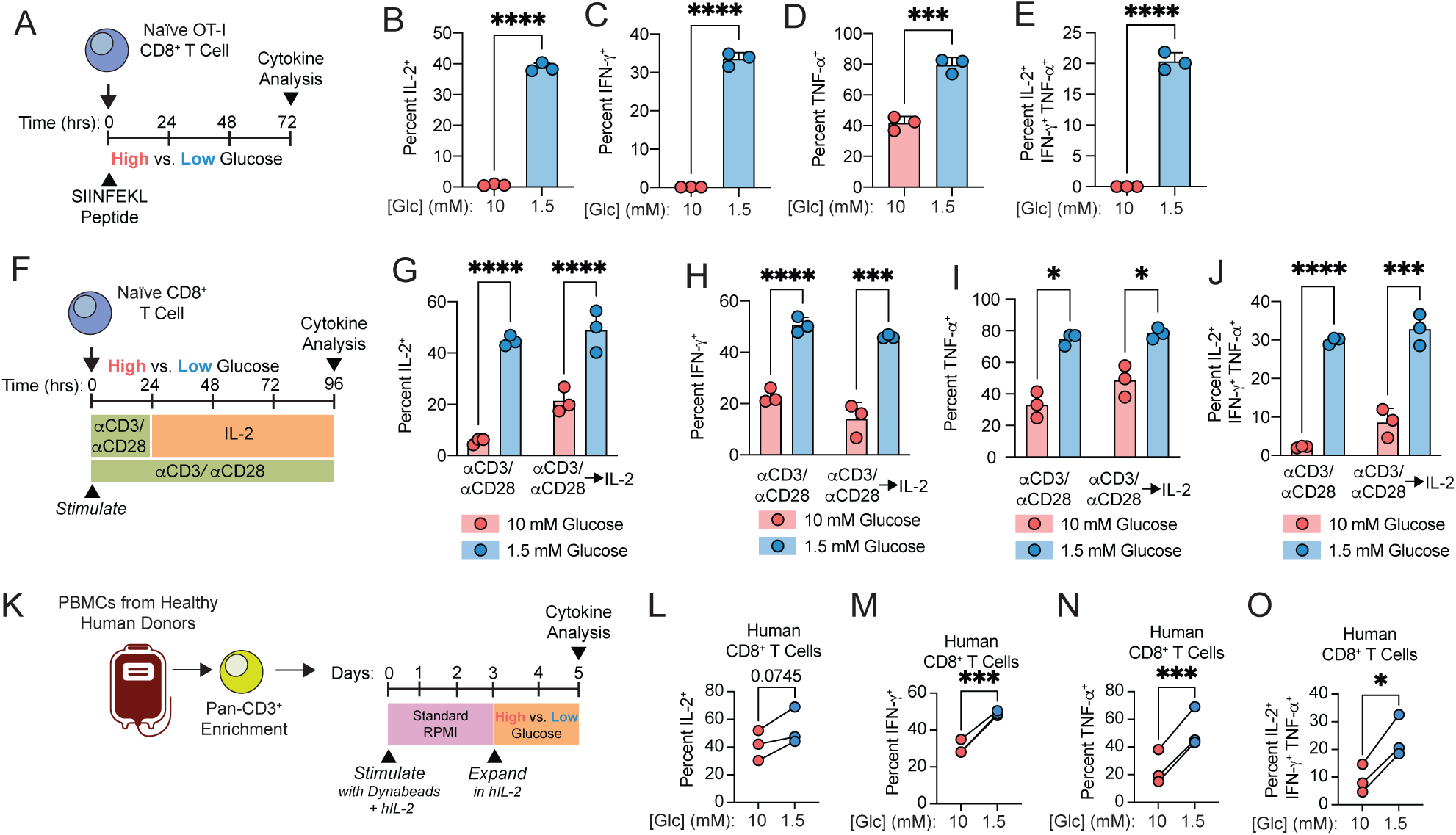
Glucose limitation raises cytokine production across activation modalities in mouse and human CD8^+^ T cells. (**A**) Experimental schematic for panels B-E. (**B-E**) Percent IL-2^+^ (**B**), IFN-γ^+^ (**C**), TNF-α^+^ (**D**), and polyfunctional (IL-2^+^ IFN-γ^+^ TNF-α^+^) (**E**) of live JAXBoy-OT-I CD8^+^ T cells after 72 hours of stimulation with SIINFEKL peptide in high (10 mM) versus low (1.5 mM) glucose. (**F**) Experimental schematic for panels G-J. (**G-J**) Percent IL-2^+^ (**G**), IFN-γ^+^ (**H**), TNF-α^+^ (**I**), and polyfunctional (IL-2^+^ IFN-γ^+^ TNF-α^+^) (**J**) of live CD8^+^ T cells after 24 hours of polyclonal activation on plate-bound α-CD3/ α-CD28 supplemented with IL-2 and then expanded in IL-2 for an additional 72 hours while maintained continuously in high (10 mM) versus low (1.5 mM) glucose. (**K**) Experimental schematic for panels L-O. (**L-O**) Percent IL-2^+^ (**L**), IFN-γ^+^ (**M**), TNF-α^+^ (**N**), and polyfunctional (IL-2^+^ IFN-γ^+^ TNF-α^+^) (**O**) of live human CD8^+^ T cells after 3 days of activation on DynaBeads in 11mM glucose supplemented with hIL-2, followed by 2 days expansion in 10 mM versus 1.5 mM glucose with hIL-2. *Abbreviations*: Glc = Glucose; hIL-2 = Human Interleukin-2; IFN-γ = Interferon Gamma; PBMCs = Peripheral Blood Mononuclear Cells; TNF-α = Tumor Necrosis Factor Alpha *Statistics*: All data points are distinct biological replicates. Data are presented as mean ± standard deviation (B-E, G-J). Panels B-E and G-J show data representative of two independent experiments where n=3 biological replicates are shown from a representative experiment. Data points in L-O are derived from three independent healthy human donors. Statistical significance assessed by Student’s t-test (B-E), two-way ANOVA (G-J), or two-tailed paired t-test (L-O). P-values: not significant (ns) for p>0.05, ∗ for p≤0.05, ∗∗ for p≤0.01, ∗∗∗ for p≤0.001, and ∗∗∗∗ for p≤0.0001.

### Glucose availability dynamically regulates polyfunctional cytokine production by T cells

The impact of glucose on cytokine expression was first evident three days post-activation, prompting us to ask whether this is an acutely adaptive response to glucose availability. After activating T cells in low glucose conditions, we raised glucose levels after either 24 or 48 hours by adding glucose to the existing cultures and then evaluated IL-2 production after 72 total hours of stimulation. Acutely raising glucose at either timepoint dampened the production of IL-2, IFN-γ, and TNF-α (**Figure 3A-D**); reduced polyfunctionality (**Figure 3E**); and increased proliferation (**Figure S3A-B**). We next titrated the amount of glucose added at 48 hours (**Figure 3F**) and found that the magnitude of decline in IL-2 (**Figure 3G**), IFN-γ (**Figure S3C**), and TNF-α (**Figure S3D**) tracked with glucose concentration. Similarly, treating activating T cells with a glucose uptake inhibitor to mimic glucose limitation within the same timeframe (**Figure 3H**) was sufficient to raise the production of IL-2 (**Figure 3I**), IFN-γ (**Figure S3E**), and TNF-α (**Figure S3F**) in a dose-responsive manner, as well as decrease proliferation (**Figure S3G**). Thus, CD8^+^ T cells adaptively raise cytokine production in response to moderate glucose limitation, and this capacity is not imprinted by metabolic exposures during early activation.

**Figure 3:**
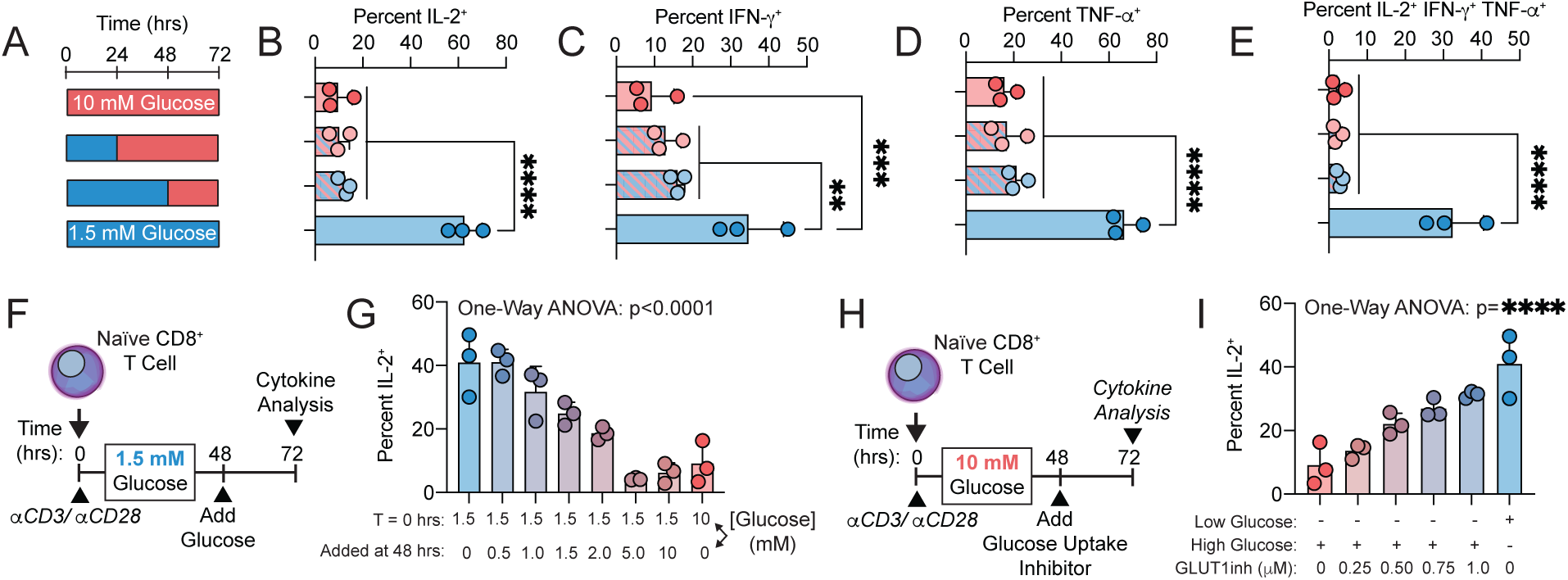
The effect of glucose availability on cytokine production is dynamic and reversible. (**A**) Experimental schematic for panels B-E. (**B-E**) Percent IL-2^+^ (**B**), IFN-γ^+^ (**C**), TNF-α^+^ (**D**), and polyfunctional (IL-2^+^ IFN-γ^+^ TNF-α^+^) (**E**) of live CD8^+^ T cells after 72 hours of activation on plate-bound α-CD3/ α-CD28. Cells were provided glucose at the indicated concentrations and times. (F) Experimental schematic for panel G. (G) Percent IL-2^+^ of live CD8^+^ T cells activated on plate-bound α-CD3/ α-CD28 in low (1.5 mM) glucose for the first 48 hours before glucose supplementation to the indicated concentrations for the last 24 hours of activation. (H) Experimental schematic for panel I. (I) Percent IL-2^+^ of live CD8^+^ T cells activated on plate-bound α-CD3/ α-CD28 at the indicated glucose concentrations. After 48 hours, a glucose uptake inhibitor (KL-11743) was added for the final 24 hours of activation. *Abbreviations*: GLUT1inh = Glucose Transporter Inhibitor KL-11743; hrs = hours; IFN-γ = Interferon Gamma; TNF-α = Tumor Necrosis Factor Alpha *Statistics*: All data points are distinct biological replicates. Data are presented as mean ± standard deviation (B-E, G, I). Panels B-E, G, and I show data representative of two independent experiments where n=3 biological replicates are shown from a representative experiment. Statistical significance assessed by one-way ANOVA (B-E, G, I). P-values: not significant (ns) for p>0.05, ∗ for p≤0.05, ∗∗ for p≤0.01, ∗∗∗ for p≤0.001, and ∗∗∗∗ for p≤0.0001.

### Citrate export by SLC25A1 limits inflammatory cytokine expression

Glucose supports T cell activation by fueling aerobic glycolysis in the cytosol and oxidative metabolism in the mitochondria. We therefore evaluated whether a shift in cytosolic versus mitochondrial metabolism controls cytokine output during moderate glucose restriction. Galactose is an alternate hexose sugar that can enter glycolysis through the Leloir pathway [46], but its catabolism generates cytosolic ATP less efficiently than glucose, causing cells to rely on oxidative phosphorylation to a greater extent [30,47–49]. Replacing glucose with galactose significantly raised IL-2 (**Figure 4A**), IFN-γ (**Figure 4B**), and TNF-α (**Figure 4C**) production. T cells activated with galactose as the carbohydrate source also displayed higher cytokine polyfunctionality (**Figure 4D**). Galactose supported proliferation to a similar extent as moderate glucose restriction (**Figure S4A**). Next, we tested whether elevated cytokine production was associated with a distinct bioenergetic state. To do this, we measured the oxygen consumption rate (OCR) as a proxy for oxidative phosphorylation and extracellular acidification rate (ECAR) as a proxy for aerobic glycolysis using a Seahorse analyzer on T cells activated in high glucose, low glucose, or galactose. Moderate glucose restriction caused small changes in baseline OCR and did not alter spare respiratory capacity (**Figures 4E-F**). Likewise, ECAR was not significantly affected when glucose was varied from 1.5 to 10 millimolar (**Figures 4G-H**). Galactose, however, markedly increased OCR and substantially reduced ECAR (**Figures 4E-H**), indicating that these T cells were performing less aerobic glycolysis and more oxidative phosphorylation. Despite displaying distinct bioenergetic profiles, both low glucose and galactose enhanced cytokine production (**Figures 4A-D**). Together, these data indicate that bioenergetic remodeling does not explain why high glucose suppresses cytokine production. As galactose causes cells to rely on oxidative phosphorylation to generate energy, we hypothesized that a metabolite derived from mitochondrial fuel oxidation regulates cytokine output. While often depicted as a closed loop within the mitochondria, intermediates from the TCA cycle are also transported to other cellular compartments. We initially focused on citrate, which can be shuttled to the cytosol by the mitochondrial citrate carrier SLC25A1 (**Figure 4I**). Once in the cytosol, citrate is converted back to acetyl-CoA that is used for lipid synthesis and protein acetylation [34,50–52]. We performed targeted polar metabolomics on activated CD8^+^ T cells cultured in high or low glucose, focusing on ratios between TCA intermediates as a measure of relative pathway organization. This revealed a significant decrease in the citrate to α-ketoglutarate ratio (**Figure 4J**) and increase in the α-ketoglutarate to malate ratio in glucose-restricted cells (**Figure 4K**), consistent with greater retention of carbon within the mitochondrial arm of the TCA cycle during glucose limitation (**Figure 4I**) [53]. On the other hand, the ratios between other TCA cycle metabolites, including citrate and malate (**Figure 4L**), as well as citrate and succinate (**Figure S4B**), were unchanged. Despite apparent alterations in the export of citrate from the mitochondria, activating CD8^+^ T cells did not secrete citrate into the media under any condition (**Figure 4M**).

**Figure 4:**
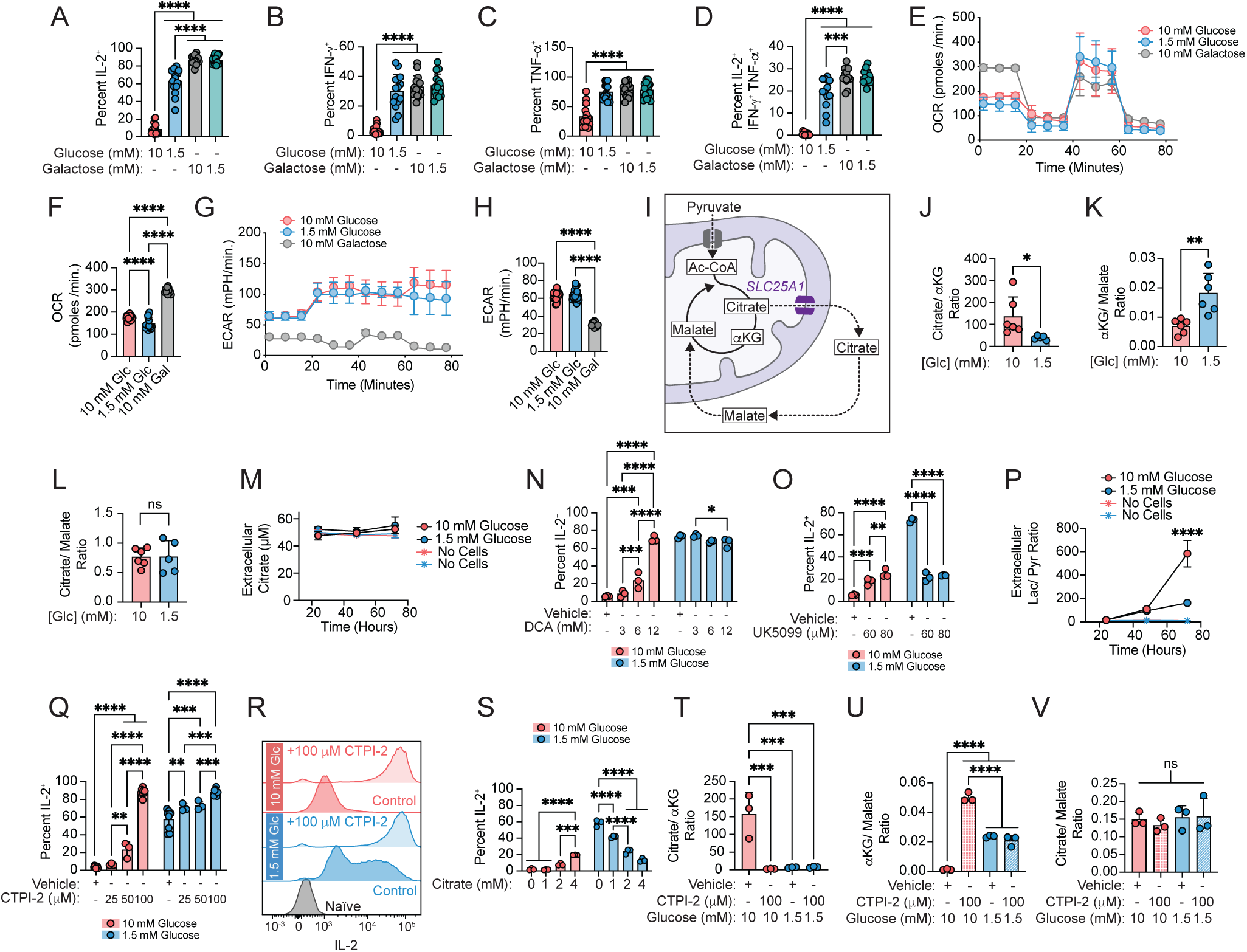
Citrate transport from the mitochondria to the cytosol by SLC25A1 limits inflammatory cytokine production. (**A-D**) Barplots depict percent IL-2^+^ (**A**), IFN-γ^+^ (**B**), TNF-α^+^ (**C**), and polyfunctional (IL-2^+^ IFN-γ^+^ TNF-α^+^) (**D**) of live CD8^+^ T cells after 72 hours of polyclonal activation on plate-bound α-CD3/ α-CD28 in glucose or galactose at the indicated concentrations. (**E-F**) Oxygen consumption rate (OCR) as measured during the Seahorse XF Cell Mito Stress Test on CD8^+^ T cells after 72 hours of polyclonal activation on plate-bound α-CD3/ α-CD28 in glucose or galactose at the indicated concentrations (**E**). Barplot depicts baseline OCR (**F**). Seahorse assay medium contained the same sugar source as the activation condition. (**G-H**) Extracellular Acidification Rate (ECAR) as measured during the Seahorse XF Cell Mito Stress Test on CD8^+^ T cells after 72 hours of polyclonal activation on plate-bound α-CD3/ α-CD28 in glucose or galactose at the indicated concentrations (**G**). Barplot depicts baseline ECAR (**H**). Seahorse assay medium contained the same sugar source as the activation condition. (**I**) Diagram depicting the fates of citrate produced in the mitochondria. (**J-L**) Metabolomics analysis on CD8^+^ T cells after 72 hours of activation on plate-bound α-CD3/ α-CD28 in high (10 mM) versus low (1.5 mM) glucose measured by liquid chromatography-mass spectrometry (LC-MS). Barplots depict ratios of peak areas for citrate to α-ketoglutarate (**J**), α-ketoglutarate to malate (**K**), or citrate to malate (**L**). (**M**) Media citrate levels by LC-MS after culturing CD8^+^ T cells for 24, 48, and 72 hours on plate-bound α-CD3/ α-CD28 in high (10 mM) versus low (1.5 mM) glucose, compared to cell-free controls. (**N**) Percent IL-2^+^ of live CD8^+^ T cells after 72 hours of polyclonal activation on plate-bound α-CD3/ α-CD28 in high (10 mM) versus low (1.5 mM) glucose with varying concentrations of dichloroacetate or vehicle control. (**O**) Percent IL-2^+^ of live CD8^+^ T cells after 72 hours of polyclonal activation on plate-bound α-CD3/ α-CD28 in high (10 mM) versus low (1.5 mM) glucose with varying concentrations of the mitochondrial pyruvate carrier (MPC) inhibitor UK-5099 or vehicle control. (**P**) Ratio of extracellular lactate to pyruvate levels measured in culture media of CD8^+^ T cells after 24, 48, and 72 hours of activation on plate-bound α-CD3/ α-CD28 in high (10 mM) versus low (1.5 mM) glucose, compared to cell-free controls. (**Q-R**) Percent IL-2^+^ of live CD8^+^ T cells after 72 hours of polyclonal activation on plate-bound α-CD3/ α-CD28 in high (10 mM) versus low (1.5 mM) glucose with varying concentrations of the mitochondrial citrate carrier inhibitor (SLC25A1) inhibitor CTPI-2 or vehicle control (**Q**). Representative flow cytometry histograms of IL-2 signal after 72 hours of activation with CTPI-2 or vehicle control, including an un-stimulated naïve control (**R**). (**S**) Percent IL-2^+^ of live CD8^+^ T cells after 72 hours of polyclonal activation on plate-bound α-CD3/ α-CD28 in high (10 mM) versus low (1.5 mM) glucose with varying concentrations of sodium citrate. (**T-V**) Metabolomics analysis on CD8^+^ T cells after 72 hours of activation on plate-bound α-CD3/ α-CD28 in high (10 mM) versus low (1.5 mM) glucose with SLC25A1 inhibition by CTPI-2. Barplots depict ratios of peak areas for citrate to α-ketoglutarate (**T**), α-ketoglutarate to malate (**U**), or citrate to malate (**V**). *Abbreviations*: Ac-CoA = Acetyl-Coenzyme A; αKG = Alpha-Ketoglutarate; CTPI-2 = Inhibitor of the Mitochondrial Citrate Carrier SLC25A1; DCA = Dichloroacetate; ECAR = Extracellular Acidification Rate; Gal = Galactose; Glc = glucose; IFN-γ = Interferon Gamma; Lac = Lactate; OCR = Oxygen Consumption Rate; Pyr = Pyruvate; TCA cycle = Tricarboxylic Acid cycle; TNF-α = Tumor Necrosis Factor Alpha; UK-5099 = Inhibitor of Mitochondrial Pyruvate Carrier *Statistics*: Data are presented as mean ± standard deviation (A-D, F, H, J-Q, S-V). Panels A-D contain data aggregated from 5 independent experiments with n=15 biological replicates. Panels E-H contain data aggregated from 3 biological replicates with 5 technical replicates each. Panels J-L contain data aggregated from 2 independent experiments with n=6 biological replicates. Panels M and P contain data aggregated from 4 biological replicates. Panels N-O, S, and T-V show data representative of two independent experiments where n=3 biological replicates are shown from a representative experiment. Panels Q-R contain data aggregated from 3 independent experiments with n=7 biological replicates. Statistical significance assessed by one-way ANOVA (A-D, F, H), Student’s t-test (J-L), or two-way ANOVA (M-Q, S-V). P-values: not significant (ns) for p>0.05, ∗ for p≤0.05, ∗∗ for p≤0.01, ∗∗∗ for p≤0.001, and ∗∗∗∗ for p≤0.0001.

Since citrate is a major TCA cycle metabolite that transits between the mitochondrial matrix and cytosol, we hypothesized that its compartmentalization regulates IL-2 production during glucose restriction. As oxidative metabolism of glucose was recently shown to retain citrate within the mitochondria [53,54], we first tested whether interventions promoting pyruvate oxidation by the TCA cycle would be sufficient to raise IL-2 production in high glucose. We therefore treated activating CD8^+^ T cells with dichloroacetate (DCA), a compound that enhances pyruvate dehydrogenase (PDH) activity inside the matrix, which commits pyruvate to mitochondrial oxidation [31,55]. DCA strongly induced IL-2 production in high glucose but had no effect in low glucose (**Figure 4N**), indicating that oxidative consumption of pyruvate potentiates IL-2 expression. By contrast, blocking pyruvate oxidation with an inhibitor of the mitochondrial pyruvate carrier (**Figure 4O**) reduced IL-2 production when T cells were cultured in low glucose. Furthermore, at later stages of activation in glucose-restricted cells, we observed a decrease in lactate excretion relative to glucose-replete conditions (**Figure 4P**), indicating that pyruvate is shuttled into the mitochondria for oxidation rather than being converted to lactate in the cytosol through aerobic glycolysis. Together, these data suggest that pyruvate oxidation favors a mitochondrial configuration that promotes IL-2 expression.

Citrate is the first TCA cycle metabolite generated from pyruvate after the PDH reaction. Following its formation in the matrix, citrate levels are regulated by mitochondrial aconitase activity, which routes citrate through the TCA cycle [54], as well as carrier-mediated transport to the cytosol by SLC25A1 [56–58]. To test the hypothesis that mitochondrial-to-cytosolic citrate transport regulates cytokine production, we treated activating CD8^+^ T cells with an inhibitor of SLC25A1 called CTPI-2. CTPI-2 potently increased IL-2 (**Figures 4Q-R**), IFN-γ (**Figure S4C**), TNF-α (**Figure S4D**), and cytokine polyfunctionality (**Figure S4E**), with a small increase in proliferation for cells cultured in low glucose (**Figure S4F**). Conversely, addition of extracellular citrate suppressed the production of IL-2 (**Figure 4S**), IFN-γ (**Figure S4G**), TNF-α (**Figure S4H**), and cytokine polyfunctionality (**Figure S4I**). Next, we considered whether citrate-dependent metabolic activities were responsible for this effect. One major fate for cytosolic citrate is its conversion into acetyl-CoA by ATP-citrate lyase (ACLY), which is used for *de novo* lipogenesis and protein acetylation [34,50–52]. However, supplementing acetate to provide an alternate substrate for cytosolic acetyl-CoA pools did not reverse the effect of SLC25A1 inhibition on cytokine production (**Figures S4J-S4M**). Treatment with the SLC25A1 inhibitor remodeled the TCA cycle to resemble glucose restriction in effector T cells (**Figures 4T-V**), reducing the citrate to α-ketoglutarate ratio (**Figure 4T**), raising the α-ketoglutarate to malate ratio (**Figure 4U**), and having no effect on the relationship between citrate and malate (**Figure 4V**). These findings implicate cytosolic citrate, and not its conversion to acetyl-CoA, in downregulating the capacity for cytokine production in effector T cells.

### Glucose availability regulates calcium-dependent transcriptional programs

We subsequently focused on IL-2 due to the large effect size of glucose availability on its production. As IL-2 is both produced and sensed by T cells [44,59], we first ruled out that IL-2 mediates its own increase during glucose limitation. Adding extra IL-2 had no impact on IL-2 production in either low or high glucose (**Figure 5A**) and did not affect proliferation (**Figure S5A**), suggesting that enhanced IL-2 production is not driven by autocrine signaling. Similarly, blocking the high affinity IL-2 receptor had little impact on IL-2 production in low glucose, although it raised IL-2 output in high glucose (**Figure 5B**), a feedback circuit that has been previously reported [23,59], and modestly increased proliferation (**Figure S5B**). To understand the underlying cellular processes, we evaluated if glucose-sensitive IL-2 production required re-stimulation. This revealed that blocking the secretory pathway for three hours in the absence of PMA and ionomycin was not sufficient to increase intracellular IL-2 levels (**Figure 5C**), indicating that most IL-2 production occurs in response to re-stimulation during that timeframe. To determine whether the increased intracellular IL-2 was actively secreted, we re-stimulated cells with PMA and ionomycin in the presence or absence of protein transport inhibitors. Under these conditions, blocking the secretory pathway markedly increased intracellular IL-2 staining (**Figure 5D**). In the absence of protein transport inhibitors, IL-2 mRNA in T cells (**Figure 5E**) and secreted IL-2 in culture medium (**Figure 5F**) was higher after 72 hours of activation under glucose restriction, consistent with increased gene expression and active IL-2 protein secretion.

**Figure 5:**
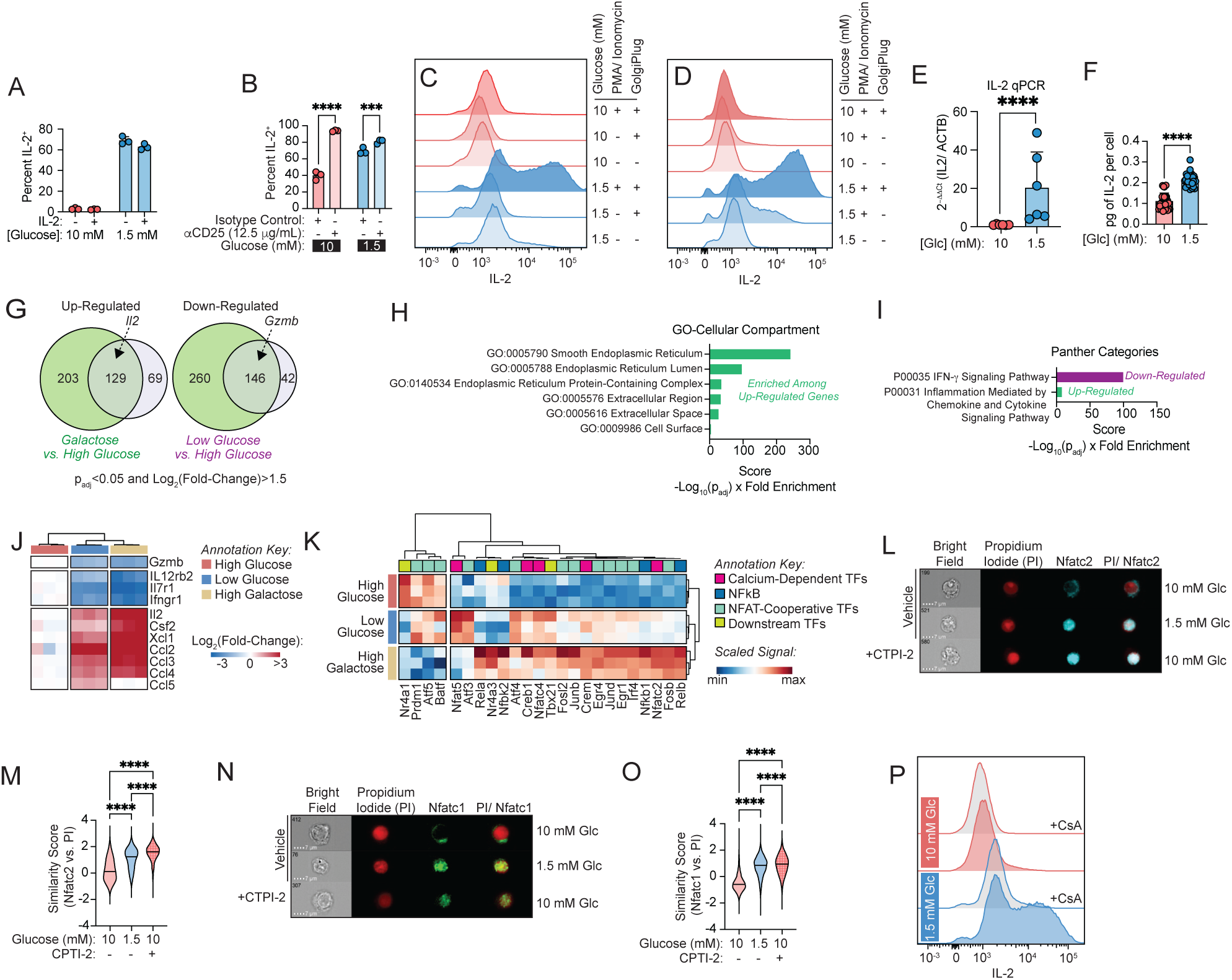
Extracellular glucose regulates calcium-dependent transcriptional programs in effector T cells. (**A**) Percent IL-2^+^ of live CD8^+^ T cells after 72 hours of polyclonal activation on plate-bound α-CD3/ α-CD28 in high (10 mM) versus low (1.5 mM) glucose. Exogenous IL-2 is supplemented at 12.5 ng/mL as indicated. (**B**) Percent IL-2^+^ of live CD8^+^ T cells after 72 hours of polyclonal activation on plate-bound α-CD3/ α-CD28 in high (10 mM) versus low (1.5 mM) glucose. Samples are treated with a CD25-blocking antibody or isotype control as indicated. (**C-D**) Flow cytometry histograms of intracellular IL-2 expression by live CD8^+^ T cells after 72 hours of activation on plate-bound α-CD3/ α-CD28 in high (10 mM) versus low (1.5 mM) glucose, where PMA/ Ionomycin and GolgiPlug are added during re-stimulation as indicated. (**E**) IL-2 mRNA expression measured by qPCR quantification in CD8^+^ cells after 72 hours of polyclonal activation on plate-bound α-CD3/ α-CD28 in high (10 mM) versus low (1.5 mM) glucose. IL-2 mRNA was normalized to actin B (ACTB) mRNA levels. (**F**) ELISA for IL-2 secretion into culture medium by CD8^+^ T cells activated on plate-bound α-CD3/ α-CD28 for 72 hours in high (10 mM) versus low (1.5 mM) glucose. (**G**) Venn diagrams of significantly up-regulated (*left*) or down-regulated (*right*) genes in bulk RNA sequencing from CD8^+^ T cells after 72 hours of polyclonal activation in galactose versus high (10 mM) glucose compared to low (1.5 mM) versus high (10 mM) glucose. (**H-I**) Gene Ontology (GO) analysis on common significantly differentially genes from analysis of bulk RNA sequencing datasets as in panel G. Up- and down-regulated genes were evaluated separately, considering only genes that were significantly differentially expressed in the same direction for galactose versus high (10 mM) glucose or low (1.5 mM) versus high (10 mM) glucose. Cellular Component (CC) gene signatures were assessed in (**H**), and Panther Pathways were assessed in (**I**). Pathways enriched among up-regulated genes are shown in green, and pathways enriched among down-regulated genes are shown in purple. (**J**) Heat map depicting the expression of cytokine genes from bulk RNA-sequencing of CD8^+^ T cells after 72 hours of polyclonal activation in the indicated conditions. (**K**) Heatmap of transcriptional programs inferred from bulk RNA-sequencing of CD8^+^ T cells after 72 hours of polyclonal activation in the indicated conditions. (**L-O**) ImageStream flow cytometry of CD8^+^ T cells after 72 hours of polyclonal activation in low (1.5 mM) versus high (10 mM) glucose, or high glucose with addition of the SLC25A1 inhibitor CTPI-2 versus vehicle control. Representative images for Nfatc2 (**L**) and Nfatc1 (**N**) are shown, as well as violin plots showing per cell similarity scores for Nfatc2 (**M**) or Nfatc1 (**O**) relative to propidium iodide (PI) nuclear signal. (**P**) Flow cytometry histograms of intracellular IL-2 expression by live CD8^+^ T cells after 72 hours of activation on plate-bound α-CD3/ α-CD28 in high (10 mM) versus low (1.5 mM) glucose, where the calcineurin inhibitor cyclosporin A is added as indicated during restimulation with PMA/ ionomycin. *Abbreviations*: ACTB = actin B; CsA = Cyclosporin A; ELISA = Enzyme-Linked Immunosorbent Assay; Glc = glucose; GO = Gene Ontology; Gzmb = Granzyme B; NFAT = Nuclear Factor of activated T cells; NFkB = Nuclear Factor Kappa-light-chain-enhancer of Activated B cells; PMA = Phorbol 12-myristate 13-acetate; qPCR = Quantitative Polymerase Chain Reaction; TFs = Transcription Factors *Statistics*: All data points shown are distinct biological replicates. Barplots are presented as mean ± standard deviation (A-B, E-F). Violin plots depict the median as a horizontal line (M, O). Panels A, B, and L-P show data representative of two independent experiments where n=3 biological replicates are shown from a representative experiment. Bulk RNA-sequencing data in panels G-K are derived from n=3 biological replicates. Panel E contains data aggregated from 2 independent experiments with n=6 biological replicates. Panel F contains data from 3 biological replicates with n=5 technical replicates, where technical replicates are shown. Statistical significance assessed by two-way ANOVA (A, B), by one-way ANOVA (F, M, O), or by Welch’s corrected two-tailed t-test (E, F). P-values: not significant (ns) for p>0.05, ∗ for p≤0.05, ∗∗ for p≤0.01, ∗∗∗ for p≤0.001, and ∗∗∗∗ for p≤0.0001.

Our observation that glucose availability regulates IL-2 gene expression prompted us to examine broader transcriptional changes by bulk RNA-sequencing. We compared effector CD8^+^ T cells activated for three days in high glucose versus low glucose or galactose, which are both conditions that induce IL-2, with the goal of defining shared transcriptional programs that could nominate a mechanism. Principal component analysis revealed clear separation among all three conditions, with the major axis of variation (explaining 66.5% of variance) distinguishing high glucose from both low glucose and galactose (**Figure S5C**). Differential expression analysis revealed 129 genes that were significantly up-regulated when comparing either low glucose or galactose to high glucose, including IL-2, as well as 146 genes that were commonly down-regulated (**Figure 5G**). Within the shared transcriptional changes, we observe an enrichment in transcripts associated with the endoplasmic reticulum (ER) by Gene Ontology (GO) analysis (**Figure 5H**), where cytokines are translated, folded, and packaged for secretion [60]. The ER also serves as a major reservoir for intracellular calcium that can be released to the cytoplasm for signaling [61,62]. Consistent with altered calcium handling, we also observe an increase in the expression of transcripts encoding for SERCA subunits, which pump calcium into the ER, and calcium chaperones that reside within the ER lumen (**Figure S5D**). Similarly, GO analysis using a curated set of PANTHER pathway categories revealed enrichment of genes associated with cytokine signaling (**Figure 5I**), with a representative subset of genes shown in **Figure 5J**. To nominate candidate upstream regulatory pathways, we inferred transcription factor activities using decoupleR [63]. Both the low glucose and galactose conditions were characterized by a prominent enrichment of calcium-responsive transcription factors (**Figure 5K**). This included members of the NFAT family of transcription factors, which are known to control IL-2 transcription in a calcium-dependent manner [64,65]. Sub-cellular localization of NFAT family members controls the expression of their target genes [66]. While phosphorylated NFAT remains in the cytoplasm, dephosphorylation by calcineurin promotes nuclear translocation and activation of NFAT-dependent genes, including IL-2 [66]. Collectively, these findings demonstrate that metabolic conditions sustaining cytokine production are characterized by activation of calcium-dependent transcription factors and ER remodeling.

To assess NFAT activation, we visualized the subcellular localization of Nfatc2 and Nfatc1 isoforms using imaging flow cytometry. Compared to cells cultured in high glucose conditions, effector CD8^+^ T cells activated under mild glucose restriction displayed increased nuclear localization of Nfatc2 (**Figures 5L-M**) and Nfatc1 (**Figures 5N-O**). Notably, inhibiting SLC25A1 with CTPI-2 in high glucose conditions recapitulated the effect of glucose restriction on NFAT nuclear localization (**Figures 5L-O**). To validate the link between calcium-signaling and IL-2 production, we treated cells with a calcineurin inhibitor during re-stimulation. Blocking calcineurin activity completely abrogated the increase in IL-2 expression observed under low glucose conditions (**Figure 5P**), indicating that NFAT mediates IL-2 production during glucose restriction. Thus, transfer of mitochondrial citrate to the cytosol serves as a metabolic checkpoint that restrains NFAT activation in effector T cells.

### Citrate restricts NFAT activation by limiting cytosolic calcium availability

Having established that mitochondrial citrate export restrains NFAT activation, we next investigated the causal mechanism. Citrate contains three carboxylate groups capable of chelating divalent cations such as calcium (**Figure 6A**). Clinically, citrate is widely used as a calcium-chelating anti-coagulant for blood collection. In mammalian physiology, citrate inhibits stone formation in the kidney by forming a soluble alternate to calcium oxalate [67], is a structural component of bone [68], and regulates bone mineral stability [68]. Using a colorimetric o-Cresolphthalein assay, we verified that citrate reduces free calcium levels in a concentration-dependent manner (**Figure 6B**). This prompted us to test whether metabolic interventions that modulate citrate production and compartmentalization affect the availability of free cytosolic calcium using Fluo-4 AM, a calcium-sensitive fluorescent dye. Consistent with the effect on NFAT localization (**Figure 5L-O**), effector cells activated in low glucose displayed higher cytosolic calcium levels compared to high glucose (**Figure 6C**). Treatment with an SLC25A1 inhibitor that retains citrate in the mitochondria significantly increased cytosolic calcium under high glucose conditions (**Figure 6C**). Additionally, adding extracellular citrate during re-stimulation phenocopied treatment with cyclosporin A (**Figures 6D-E**). As cyclosporin A blocks the NFAT pathway, these results suggest that citrate is sufficient to suppress calcium-dependent NFAT activation.

**Figure 6:**
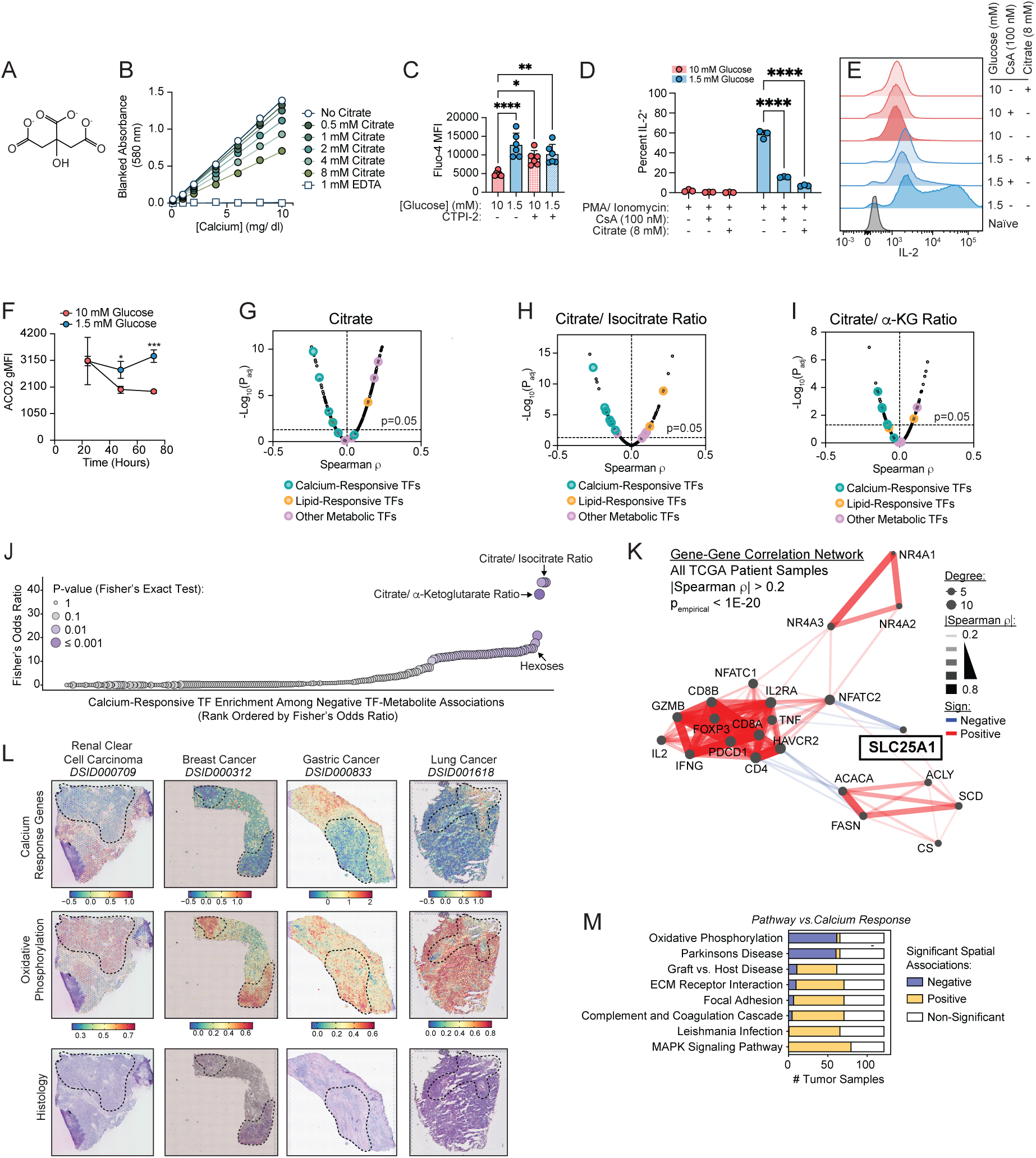
Cytosolic citrate reduces free calcium and limits calcium-dependent transcriptional programs. (**A**) Chemical structure of citrate showing three carboxylate groups that can chelate Ca^2+^. (**B**) Blanked absorbance of o-Cresolphthalein bound to calcium with increasing concentrations of citrate or EDTA, across a range of calcium concentrations. (C) Mean Fluorescence Intensity (MFI) of Fluo-4 dye in live CD8^+^ T cells after 72 hours of polyclonal activation on plate-bound α-CD3/ α-CD28 in high (10 mM) versus low (1.5 mM) glucose with addition of the SLC25A1 inhibitor CTPI-2 or vehicle control. (**D-E**) Percent IL-2^+^ of live CD8^+^ T cells after 72 hours of polyclonal activation on plate-bound α-CD3/ α-CD28 in high (10 mM) versus low (1.5 mM) glucose. The calcineurin inhibitor cyclosporin A or sodium citrate are added during re-stimulation with PMA/ ionomycin as indicated. Barplots shown in (**D**) and representative flow cytometry histograms shown in (**E**). (**F**) Dotplots depicting geometric Mean Fluorescence Intensity (gMFI) of mitochondrial aconitase 2 (ACO2) by intracellular flow cytometry staining in live CD8^+^ T cells across 72 hours of polyclonal activation by plate-bound α-CD3/ α-CD28 in high (10 mM) versus low (1.5 mM) glucose. (**G-I**) Positive and negative correlations between transcription factor programs calculated with decoupleR and selected metabolite levels: citrate (**G**), citrate/ isocitrate ratio (**H**), and citrate/ α-Ketoglutarate ratio (**I**) across cancer cell lines from the Cancer Cell Line Encyclopedia (CCLE). (**J**) Calcium-responsive transcription factor programs ranked by enrichment among all transcription factor programs with significant negative associations between program score and metabolite abundance across cancer cell lines in the Cancer Cell Line Encyclopedia (CCLE). (**K**) Gene-gene correlation network across TCGA tumors (n=9,279 patient tumor samples). Pairwise Spearman correlations were calculated from bulk tumor RNA-sequencing data. Nodes represent genes involved in lipid metabolism, calcium signaling, and T cell function. Edges connect gene pairs exhibiting significant correlations (|ρ| > 0.2, p-value<1E-20). Edge color indicates the direction of the correlation, and edge width is proportional to correlation strength. Node size is proportional to network degree. (**L**) Representative images of Visium human tumor tissue sections overlaid with the calcium response gene signature score or KEGG oxidative phosphorylation gene signature score. Regions with low or high scores are manually annotated for visual clarity. (**M**) Barplots depicting the number of Visium tumor samples with significant cross-correlation (FDR-adjusted p-value < 0.001 and |Moran’s I| > 0.1) between the indicated KEGG pathway and the calcium response signature score. *Abbreviations*: ACO2 = Aconitase 2; CCLE = Cancer Cell Line Encyclopedia; CsA = Cyclosporin A; EDTA = ethylenediaminetetraacetic acid; FDR = False Discovery Rate; gMFI = Geometric Mean Fluorescence Intensity; HIF1A = Hypoxia-Inducible Factor 1-Alpha; Lac = Lactate; MFI = Mean Fluorescence Intensity; PMA = Phorbol 12-myristate 13-acetate; Pyr = pyruvate; TCGA = The Cancer Genome Atlas; TFs = Transcription Factors *Statistics*: All data points shown are distinct biological replicates. All plots depict mean ± standard deviation (C, D, F). Panel C contains aggregated from 2 independent experiments with n=6 biological replicates. Panels D-E show data representative of two independent experiments where n=3 biological replicates are shown from a representative experiment. Panel F shows data averaged from four biological replicates. Statistical significance assessed by on-way ANOVA (C) or two-way ANOVA (D). All other statistics are detailed in the Materials and Methods. P-values: not significant (ns) for p>0.05, ∗ for p≤0.05, ∗∗ for p≤0.01, ∗∗∗ for p≤0.001, and ∗∗∗∗ for p≤0.0001.

Several metabolic enzymes and transporters are known to regulate citrate partitioning between mitochondrial and cytosolic compartments. In addition to the citrate carrier (SLC25A1), mitochondrial aconitase (ACO2) commits citrate to mitochondrial oxidation through an equilibrium reaction that is largely independent of citrate concentration [54,69]. We therefore evaluated how ACO2 expression changes over three days of activation in low versus high glucose. During early activation, when T cells express high IL-2 regardless of glucose condition, ACO2 was also expressed at consistently high levels (**Figure 6F**). After three days, however, cells activated in low glucose maintained higher ACO2 expression than counterparts activated in high glucose (**Figure 6F**). Given that ACO2 has been previously shown to promote mitochondrial oxidation of citrate [70], this suggests that sustaining high ACO2 expression may maintain NFAT-dependent transcription during the effector phase.

Although NFAT is particularly important for T cell activation and function, calcium is a ubiquitous second messenger in mammalian cells, raising the question of whether citrate compartmentalization regulates calcium-dependent processes more broadly in mammalian biology. To test this, we asked whether signatures of other calcium-dependent transcriptional programs statistically associate with citrate-related metabolites across a large panel of cancer cell lines representing a wide range of tissue types and oncogenic drivers. We curated cancer cell lines from the Cancer Cell Line Encyclopedia (CCLE) with matched transcriptomic and metabolomic datasets [32,71]. After inferring transcription factor activities for each cell line, we correlated these values with intracellular levels of TCA cycle metabolites as well as metabolite ratios that report on partitioning between citrate export and mitochondrial oxidation. As validation for this approach, we observed a strong positive association between lactate levels and HIF1A transcriptional activity, a known regulator of glycolytic gene expression [72–74] (**Figure S6A**). Additionally, consistent with a role for citrate as a substrate for lipogenesis, we recovered positive associations between citrate and the citrate/ isocitrate ratio with SREBP transcriptional programs (**Figures 6G-H**). We next focused on calcium-responsive transcriptional programs. This analysis revealed significant negative associations between calcium-responsive transcription factor programs and citrate (**Figure 6G**), the citrate/ isocitrate ratio (**Figure 6H**), and citrate/ α-Ketoglutarate ratio (**Figure 6I**). As single metabolites, however, α-Ketoglutarate and isocitrate both displayed fewer significant associations with calcium-responsive transcription programs (**Figures S6B-C**). To assess this pattern in a more systematic manner, we evaluated for each metabolite whether calcium-responsive transcription factors (MEF2A, MEF2B, MEF2C, NFATC2, NFATC1, and CREM) were overrepresented among transcriptional programs with significant negative correlations. From this analysis, the citrate/ α-Ketoglutarate and citrate/ isocitrate ratios displayed some of the highest Fisher’s odds ratios out of all metabolites, denoting a greater likelihood of significant negative associations with calcium-responsive transcriptional programs (**Figure 6J**). Hexoses were also among the top-ranked features, consistent with high glucose limiting NFAT activity in effector T cells (**Figure 6J**). Collectively, these findings suggest that citrate compartmentalization is a general cellular mechanism that regulates calcium-dependent processes in the cytosol.

SLC25A1 has been largely studied as a component of the lipogenic pathway, since cytosolic citrate serves as a major substrate for fatty acid synthesis [75–78]. Having identified a new role for SLC25A1 in regulating calcium-dependent programs in T cells and cancer cell lines, we asked whether evidence for this function could also be detected in human tumors. We therefore examined correlations between SLC25A1 gene expression with transcripts encoding lipogenic machinery, immune cell markers, and calcium-responsive transcription factors across nearly 10,000 human patient tumor samples in The Cancer Genome Atlas (TCGA) [79]. Consistent with the established role for SLC25A1 in lipogenesis, we recover significant positive associations between SLC25A1 and lipogenic enzymes (**Fig. 6K**). As expected, we also find significant positive associations between NFATc2 (an NFAT family transcription factor) with calcium-dependent nuclear receptors, as well as between NFATc2 and T cell lineage genes (**Fig. 6K**). Using the same analysis, we identify a significant negative association between SLC25A1 and NFATc2 (**Fig. 6K**). Thus, SLC25A1 sits at the nexus between *de novo* lipogenesis and calcium-dependent immune programs in human tumors.

Having uncovered a functional connection between calcium-dependent transcriptional states and oxidative metabolism in mammalian cells, we hypothesized that spatial variation in calcium signaling within tissues should be associated with distinct mitochondrial phenotypes. To test this, we assembled 122 Visium spatial transcriptomic datasets from human patient tumors (**Table 1**) and systematically evaluated spatial cross-correlations between a calcium response gene signature and KEGG pathways [80]. This enabled us to define multicellular tissue neighborhoods and evaluate how KEGG pathway activity was distributed around spots with high or low calcium-response scores. Regions exhibiting low calcium-responsive scores tended to be spatially adjacent to spots with high expression of oxidative phosphorylation genes across all tumor types (**Fig. 6L**). Moreover, oxidative phosphorylation was the top KEGG pathway showing significant negative enrichment in relation to calcium response programs across the greatest number of tumors (**Fig. 6M**). This provides further evidence that mitochondrial citrate clearance sustains the activity of calcium-dependent processes by limiting cytosolic citrate that suppresses free calcium.

## Discussion

Here, we have uncovered a regulatory mechanism by which mitochondria control cytosolic calcium signaling through the subcellular distribution of citrate. Calcium signaling is an absolute requirement for effector T cell activation and acquisition of inflammatory functions. First implicated in lymphocyte biology over a half century ago [81,82], calcium has emerged as a major second messenger induced directly downstream of T cell stimulation [83,84]. Up to 75% of the transcriptional changes in activating T cells are now known to depend on calcium signaling [85]. Much of this is driven by the NFAT family of transcription factors, which require calcium-dependent dephosphorylation by calcineurin to enter the nucleus [66], where they coordinate an inflammatory gene expression program [64,86]. As a result, cytosolic calcium must be tightly regulated. A network of ion channels, transporters, pumps, and buffering proteins control calcium dynamics [83,87,88], and our work now identifies mitochondrial citrate oxidation as an additional layer of control over this process.

In CD8^+^ T cells, we find that extracellular glucose availability dictates the cellular fate of citrate and, consequently, calcium-dependent transcriptional programs. When glucose is limited, T cells favor mitochondrial citrate oxidation over cytosolic export (**Figure 4J-K**). Since citrate can chelate calcium, its retention in the mitochondria raises the availability of cytosolic calcium and sustains nuclear localization of NFAT. This pathway is acutely responsive to changes in extracellular glucose (**Figure 5L-O**), suggesting that entry into low-glucose environments such as hypoxic niches and tumors may dynamically reprogram the capacity for cytokine production by effector T cells through this mechanism. We also found that raising extracellular citrate is sufficient to dampen cytosolic calcium signaling during glucose restriction (**Figure 4S**). It is still unclear which transporter mediates citrate uptake by T cells, but an isoform of SLC25A1 has been reported to localize to the plasma membrane that may serve this purpose [89]. Suppressing cytokine production required treatment with millimolar concentrations of citrate, which is an order of magnitude higher than plasma concentrations [90,91]. However, there are specialized niches for handling citrate in the body where citrate exists at much higher levels. Bone serves as a major citrate reservoir [92–97], while tissues such as the kidney and prostate are known to take up and secrete high quantities of citrate [91,98–100]. For example, expressed prostate secretion, the liquid component of seminal fluid, contains 50-80 mM citrate [100–104]. How these different citrate environments impact functional specialization among immune cells and other calcium-dependent pathways remains an open question.

We also identify the mitochondrial citrate carrier as a major cellular node that regulates cytosolic free calcium. However, there are other pathways and enzymes that impact citrate levels and compartmentalization, raising the possibility that calcium regulation by mitochondrial-derived citrate is connected to a wider range of additional processes that were previously unappreciated. Mitochondrial ACO2 is the enzyme that consumes citrate through its conversion into isocitrate as part of the TCA cycle and plays a significant role in cellular citrate clearance [70]. Since ACO2 catalyzes a near-equilibrium reaction [54,105,106], movement through the TCA cycle is sensitive to aconitase activity and expression levels, as well as downstream reactions that consume isocitrate, which collectively pull the net cycle forward. For example, zinc is a natural inhibitor of mitochondrial aconitase [107,108] and the mitochondrial zinc transporter is highly expressed by mammalian cells that secrete citrate to disable aconitase activity [109–111]. Calcium itself also regulates mitochondrial processes that impact citrate. Raising mitochondrial calcium enhances matrix dehydrogenase activity [112–114], amplifying the entry of pyruvate into the TCA cycle as well as promoting citrate oxidation. Thus, our work suggests that pathways upstream of mitochondrial citrate metabolism may functionally regulate calcium-dependent signaling programs.

With the advent of adoptive T cell therapies for cancer treatment, the ability to modulate NFAT activity has clear translational implications. During cell manufacturing, our findings suggest that adjusting glucose or citrate concentrations, or genetically targeting SLC25A1 or ACO2, could be used to bias the representation of cell states in the final product and to prevent reprogramming upon entering the tumor microenvironment. Targeting metabolic pathways that regulate core T cell programs may offer a means to tune T cell function without disabling essential pathways.

Cytokine production in effector T cells is a particularly strong read-out of calcium signaling, because it is potently driven by NFAT family transcription factors. However, calcium is a ubiquitous second messenger in mammalian biology that coordinates a much wider range of cellular processes. For example, in addition to its role in T lymphocytes, the NFAT family member NFATc1 contributes to VEGF-driven angiogenic programs in endothelial cells [115–117], and our work implies that this process may be subject to metabolic regulation. Our analysis across hundreds of human cancer cell lines reveals that citrate broadly regulates calcium-dependent transcription factors (**Figure 6G-K**), which has many potential implications. As an example, many leukemic cells rely on calcium-sensitive MEF2 family transcription factors for proliferation and survival [118–120], which may render them vulnerable to interventions disrupting citrate metabolism or compartmentalization. In addition to transcription, calcium is known to regulate a wide array of cellular processes such as vesicular trafficking [121,122], cytoskeletal dynamics [123–125], muscle contraction [126,127], mitochondrial dehydrogenase activity [112–114], cell cycle progression [128,129], and apoptosis [130,131]. By extension, any cellular condition that remodels citrate partitioning between mitochondrial oxidation and cytosolic export may impact these kinds of programs that depend on calcium signaling, fundamentally connecting mitochondrial activity and fuel availability to cellular behavior.

### Limitations

Our data support a causal link where mitochondrial citrate export reduces free cytosolic calcium that is consistent with a direct chelating mechanism. However, it is important to note that citrate also serves as a precursor for cytosolic acetyl-CoA that feeds lipogenesis and is used as a donor for protein acetylation [34,50–52]. This work does not exclude other cellular fates for citrate that may still be important in the effector phase of a T cell response. Indeed, citrate may serve multiple roles concurrently in effector T cells, and it will be important for future studies to reveal how partitioning between competing processes regulates T cell state and function. Another open question relates to the temporal regulation of this process, specifically why the effect of citrate on cytokine production only becomes apparent three days into the effector response. Lastly, our present studies focus on a particular window in T cell activation, capturing the transition from a naïve cell through clonal expansion and into an effector state. In tumors, where persistent antigen stimulation, metabolic dysfunction, and hypoxia drive progressive T cell dysfunction and exhaustion, this regulatory pathway may operate differently than it does during effector differentiation and will be an important area for future investigation.

## Methods

### Resource Availability

#### Lead contact

Further information and requests for resources should be directed to and will be fulfilled by the Lead Contact, Alison E. Ringel.

#### Materials availability

No new unique materials were generated in this study.

#### Data and code availability

Raw RNA-sequencing data files will be made available in the NCBI GEO repository upon publication. All code will be made available by the lead author upon request.

### Experimental Model and Subject Details

#### Mice

All mouse colonies and experimental animals were housed within the HPPF barrier facility at the Ragon Institute of Mass General, MIT, and Harvard. All animals were housed with free access to food and water, controlled temperature, and a 12-hour light-dark cycle. All animal experiments were approved by the Institutional Animal Care and Use Committee (IACUC) of Massachusetts General Hospital (MGH) under the Animal Study Protocol 2022N000056. The MGH Center for Comparative Medicine (CCM) is accredited by the Association for Assessment and Accreditation of Laboratory Animal Care (AAALAC) International. For experiments with wild-type mice, cohorts of 8 to 12-week-old female C57BL/6J (strain #000664) were purchased from Jackson Laboratories. OT-I mice (C57BL/6-Tg(TcraTcrb)1100Mjb/J, strain #003831) and congenic CD45.1/CD45.1 JAXBoy mice (C57BL/6J-*Ptprc^em6Lutzy^*/J, strain #033076) were purchased from Jackson Laboratories. These animals were crossed in-house to create a CD45.1/CD45.2 JAXBoy-OT-I strain. All animals were used for experiments between 10-16 weeks of age.

#### Primary human immune cells

Primary blood mononuclear cells (PBMCs) were isolated from buffy coats via centrifugation with Lymphoprep (Stem Cell Technologies, 07811) in SepMate PBMC isolation tubes (Stem Cell Technologies, 85450). Remaining contaminating red blood cells were lysed with ACK lysing buffer (Gibco, A1049201) for 3 minutes at room temperature. Untouched CD3^+^ cells were isolated using the EasySep Human CD3 Negative Selection Kit (Stem Cell Technologies, 17951) according to manufacturer instructions. CD3^+^ T cells were stimulated with anti-CD3/CD28 DynaBeads (ThermoFisher, 11161D) at a 1:1 cell: bead ratio for three days in standard RPMI (11 mM glucose) supplemented with 10% FBS, 1% penicillin-streptomycin, 1 mM sodium pyruvate, 10 mM HEPES, and 10 ng/mL human IL-2 (PeproTech, 200-02-50G). Cells were magnetically removed from DynaBeads after three days, and re-cultured in RPMI with the indicated glucose concentration and fresh IL-2 for two more days.

### Method Details

#### Primary murine T cell isolation and *ex vivo* culture

Primary T cells were isolated after harvesting and pooling the spleen, inguinal lymph nodes, and axillary lymph nodes from 10 to 16-week-old mice. Tissues were mechanically disrupted, passed through a 70-micron filter, and washed in MACS buffer (1% heat inactivated-FBS and 2 mM EDTA) before lysing red blood cells with ACK lysing buffer (Gibco, A1049201) for 2 minutes at room temperature. Naïve CD8^+^ T cells were isolated using negative selection via the EasySep^TM^ Mouse Naïve CD8^+^ T cell Isolation Kit (Stem Cell Technologies, 19858). Where indicated, naïve cells were stained with Cell Trace Violet (Thermo Scientific, C34557) according to manufacturer instructions. 100,000 cells were seeded in flat-bottom 96-well plates pre-coated with 4 μg/mL each of α-CD3 (Bio X Cell, BE0001-1) and α-CD28 (Bio X Cell, BE0015-1) antibodies. Where indicated, JAXBoy-OT-I T cells were alternatively activated with 1 µg/mL of SIINFEKL peptide derived from hen egg white ovalbumin corresponding to amino acids 257-264 (AnaSpec, AS-60193-1) and 12.5 ng/mL of murine IL-2 (Miltenyi Biotec, 130-120-330). Naïve cells were maintained as controls in each assay with 10 ng/mL of murine IL-7 (Peprotech, 217-17). For experiments that analyzed cytokine production, T cells were stimulated with 10 ng/mL Phorbol 12-myristate 13-acetate (PMA) (Millipore-Sigma, P8139) and 500 ng/mL ionomycin (Millipore-Sigma, I0634) for an additional 3 hours at 37°C in the presence of 1 µg/mL BD GolgiPlug^TM^ (BD Biosciences, 555029). In some cytokine experiments, 100 nM cyclosporin A (MedChemExpress, HY-B0579) was added during re-stimulation. CTPI-2 (MedChemExpress, HY-123986) was diluted in DMSO and used in cell culture across concentrations ranging from 25 to 100 µM. Trisodium citrate dihydrate (Fisher Scientific, AA3643936) was diluted in sterile water and used in cell culture across concentrations ranging from 1 to 8 mM. An inhibitor of glucose uptake via GLUT1, KL-11743 (MedChemExpress, HY-145597) was diluted in DMSO and used in cell culture across concentrations ranging from 0.25 µM to 1 µM. An inhibitor of the mitochondrial pyruvate carrier (MPC), UK5099 (MedChemExpress, HY-15475), was diluted in DMSO and used in cell culture at 60 and 80 µM. An inhibitor of pyruvate dehydrogenase kinase (PDK), dichloroacetate (MedChemExpress, HY-Y0445A), was diluted in water and used in cell culture at 3, 6, 12 µM. Sodium acetate (ThermoFisher, AAA1318430) was dissolved in PBS and used in cell culture at a concentration of 7.5 mM. The *InVivo*Mab anti-mouse CD25 blocking antibody clone PC-61.5.3 (Bio X Cell, BE0012) was used in cell culture at 12.5 µg/mL, and the corresponding isotype control *InVivo*Mab rat IgG1 anti-horseradish peroxidase (Bio X Cell, BE0088) was used at the same concentration. Galactose (ThermoFisher, AAA1281330) was supplemented to media in some experiments, and used at a final concentration of 1.5 mM or 10 mM. Cells were incubated at 37°C in humidified 5% CO2 incubator for the duration of each experiment. Media consisted of glucose-free RPMI (ThermoFisher, 11879020) supplemented with 10% FBS, 1 % penicillin-streptomycin (ThermoFisher, 15140-122), 1 mM sodium pyruvate (ThermoFisher, 11-360-070), 10 mM HEPES (ThermoFisher, 15-630-080), and 110 µM ß-mercaptoethanol (ThermoFisher, 21985023). Sterile glucose solution (ThermoFisher, A2494001) was added to the above-prepared glucose-free RPMI to reach the indicated concentrations. All three mouse strains were used interchangeably for polyclonal T cell activation experiments, while ovalbumin peptide stimulation experiments utilized T cells derived from JAXBoy-OT-I mice.

#### Primary human T cell isolation and *ex vivo* activation

Primary blood mononuclear cells (PBMCs) were isolated from leukopaks via centrifugation with Lymphoprep (Stem Cell Technologies, 07811). Red blood cells were lysed with ACK lysing buffer (Gibco, A1049201) for 5 minutes at room temperature. Pan CD3^+^ cells were isolated using the EasySep Human CD3 Positive Selection Kit (Stem Cell Technologies, 17851) according to manufacturer instructions. CD3^+^ T cells were stimulated with DynaBeads (ThermoFisher, 11161D) at a 1:1 ratio for three days in standard RPMI (11 mM glucose) supplemented with 10% FBS, 1% penicillin-streptomycin, 1 mM sodium pyruvate, 10 mM HEPES, and 10ng/mL human IL-2 (PeproTech, 200-02-50G). Cells were removed from DynaBeads after three days, and re-cultured in RPMI with the indicated glucose concentration and fresh IL-2 for two more days.

#### Antibody staining and flow cytometry

Mouse cells were first stained with LIVE/DEAD Fixable Near-IR stain (ThermoFisher Scientific, L10119) diluted in 1X DPBS (ThermoFisher, 14190-250) according to manufacturer’s instructions. Human cells were first stained with Zombie Violet™ Fixable Viability Kit (BioLegend, 423113) according to the manufacturer’s instructions. Extracellular antibody staining was performed in MACS buffer (1X DPBS containing 1% FBS and 2 mM EDTA) for 30 minutes at 4°C. Intracellular staining was performed according to manufacturer’s instructions using the Cytofix/Cytoperm Fixation/Permeabilization Solution Kit (BD Biosciences, 555028). Primary antibodies for extracellular markers were used at a 1:200 dilution, while antibodies for intracellular antigens were used at a 1:100 dilution. For unconjugated antibodies, secondary antibodies were used in a secondary stain at a 1:1000 dilution. Data were analyzed using FlowJo version 10.10.1.

The following antibodies were used for staining mouse cells for flow cytometry: Anti-aconitase 2 antibody [6F12BD9] (Abcam, ab110321), Alexa Fluor™ 488 goat anti-mouse IgG1 cross-adsorbed secondary antibody (Invitrogen, A-21121), PE anti-mouse IL-2 antibody (BioLegend, 503808), PE/Cyanine7 anti-mouse CD69 antibody (BioLegend, 104512), Brilliant Violet 605™ anti-mouse CD62L antibody (BioLegend, 104437), PerCP/Cyanine5.5 anti-mouse IFN-γ antibody (BioLegend, 505822), PE/Cyanine7 anti-mouse IFN-γ Antibody (BioLegend, 505825), Brilliant Violet 650™ anti-mouse TNF-α antibody (BioLegend, 506333), PerCP/Cyanine5.5 anti-mouse TNF-α Antibody (BioLegend, 506321), BD Horizon™ BUV395 rat anti-mouse CD25 (BD Biosciences, 564022), Brilliant Violet 510™ anti-mouse CD69 antibody (BioLegend, 104531), Brilliant Violet 650™ anti-mouse CD69 antibody [H1.2F3] (BioLegend, 104541), and FITC anti-mouse TNF-α Antibody (BioLegend, 506304), Goat anti-Mouse IgG1 Cross-Adsorbed Secondary Antibody, Alexa Fluor™ 350 (ThermoFisher, A21120).

The following antibodies were used for staining human cells for flow cytometry: FITC anti-human CD69 antibody (BioLegend, 310904), PE/Cyanine7 anti-human CD45RA antibody (BioLegend, 304126),

Brilliant Violet 605™ anti-human CD4 antibody (BioLegend, 317438), APC anti-human CD8α antibody (BioLegend, 301014), PerCP/Cyanine5.5 anti-human TNF-α antibody (BioLegend, 502926), PE anti-human IFN-γ antibody (BioLegend, 502510), and Brilliant Violet 650™ anti-human IL-2 Antibody (BioLegend, 500334).

#### Enzyme-linked immunosorbent assays (ELISA)

Media supernatants were collected, spun at 300 *g* for 5 minutes to remove remaining cells, and stored at -20°C. An ELISA assay was performed according to manufacturer’s instructions for murine IL-2 (Biolegend, 431001). ELISA assays were read on a Synergy Neo2 plate reader (BioTek) and normalized to live cell counts per well.

#### Intracellular calcium assay

Activated T cells were loaded with 3 μM Fluo-4 dye (ThermoFisher, F14201) in RPMI with 2.5 mM probenecid (Invitrogen, P36400) and incubated at 37°C for 30 minutes. RPMI used in this assay was supplemented with glucose to match the glucose concentrations that were present during activation in each condition. Cells were then washed twice and resuspended in dye-free RPMI for another 20 minutes at 37°C to allow full esterification of AM-esters. Cells were resuspended in 1X DPBS containing calcium and magnesium (ThermoFisher, 14040133) and analyzed via flow cytometry to measure free intracellular calcium.

#### Extracellular calcium quantification assay

The Calcium Assay Kit (Cayman Chemical, 701220) was used according to kit instructions to quantify the calcium-chelating ability of trisodium citrate dihydrate (Fisher Scientific, AA3643936) across a range of concentrations. Ethylenediaminetetraacetic acid (EDTA) (ThermoFisher, 15575020), a high-affinity calcium chelator, was used as a positive control.

#### RNA extraction and RT-qPCR

Cytokine gene expression analysis was quantified using reverse transcriptase quantitative PCR (RT-qPCR). Bulk RNA was extracted from approximately 1.5 million T cells per condition and purified from cells using the Direct-zol RNA Miniprep Kit (Zymo Research, R2053). RT-qPCR was performed using the iTaq Universal SYBR Green One-Step Kit (Bio-Rad, 1725151) on a QuantStudio 3 PCR System and analyzed using ΔΔC_t_ calculations. β-actin was used as a reference gene for normalization.

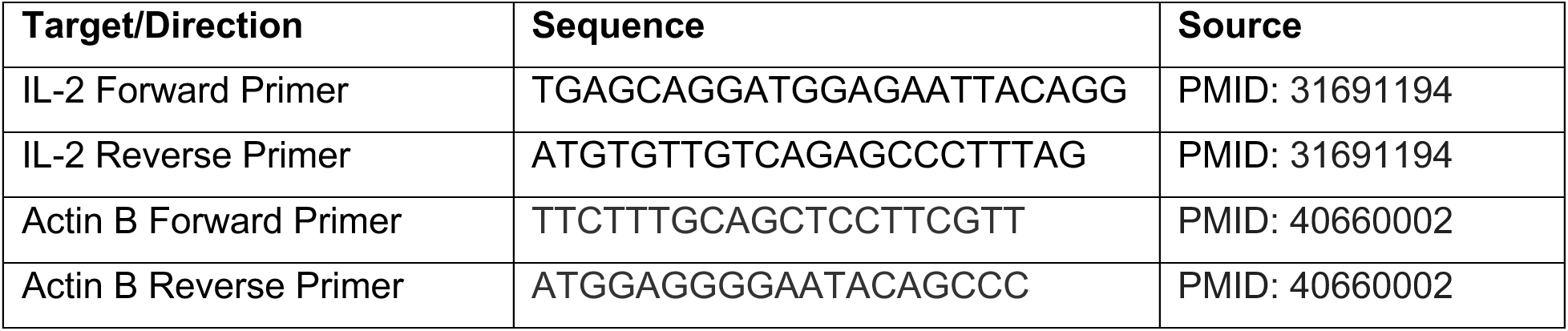

#### Bulk RNA-sequencing

RNA was preserved from approximately 100,000 activated T cells per condition in 50 μL of DNA/RNA Shield (Zymo Research, R1100). Bulk RNA-sequencing was performed by Plasmidsaurus.

#### Extracellular flux analysis

Extracellular acidification and oxygen consumption were measured using a Seahorse XF Pro Analyzer (Agilent Technologies). XF Pro Cell Culture Microplates were coated with 50 μg/mL Poly-D-lysine (Sigma-Aldrich, A-003-E) and allowed to dry before use. Seahorse FluxPaks were hydrated and used according to the manufacturer’s instructions (Agilent Technologies, 103793-100). T cells were pre-activated for 72 hours in the indicated amount of glucose or galactose and resuspended in XF RPMI Seahorse medium (Agilent Technologies, 103576-100) containing a matched amount of glucose or galactose and 1 mM pyruvate, and 2 mM glutamine. On the day of the assay, 150,000 cells were spin-seeded per well and the Seahorse Cell Mito Stress test was performed according to the manufacturer’s instructions using 1 μM oligomycin A, 1 μM FCCP, and a combination of 0.5 μM antimycin A with 0.5 μM rotenone (Agilent Technologies, 103015-100). The resulting datasets were analyzed in Wave Desktop software (version 2.6).

#### Imaging flow cytometry analysis

Activated T cells were stained with LIVE/DEAD Fixable Near-IR stain (Thermo Scientific, L10119) and fixed/permeabilized with the Foxp3/Transcription Factor Fixation/Permeabilization Kit (ThermoFisher, 00-5521-00) according to manufacturer instructions. Cells were stained intracellularly with Alexa Fluor® 647 NFAT1 (D43B1) rabbit monoclonal antibody (Cell Signaling Technology, 14201S) at 1:100 dilution, Alexa Fluor® 488 anti-NFATc1 antibody (BioLegend, 649604) at 1:100 dilution, and 1 μg/mL of propidium iodide nuclear stain (Fisher Scientific, ICN19545825) in 1X permeabilization buffer at room temperature for 30 minutes. Samples were imaged on the ImageStreamX Mark II imaging cytometer. The Nuclear Localization Wizard was utilized in the IDEAS software application (version 6.4) to quantify nuclear localization of the Nfatc2 and Nfatc1 probes. Cells were first gated to exclude debris and dead cells, followed by selecting cells with double positive signal for nuclear and NFAT stains.

#### Intracellular metabolite extraction and analysis via GC/MS

Activated T cells were washed with 0.9% NaCl on ice, and metabolites were extracted with 80% cold methanol containing 1 µM ^13^C/^15^N-labeled Canonical Amino Acid Mix (Cambridge Isotope Laboratories, MSK-CAA-1) as an internal standard. Cells were vortexed vigorously for 10 minutes at 4°C, before pelleting to remove debris and then dried down under nitrogen gas. Metabolites were derivatized as previously described [132]. Briefly, metabolites were incubated with 16 µl MOX reagent (ThermoFisher, TS-45950) at 37°C for 1 hour followed by addition of 20 µl of N-(tert-Butyldimethylsilyl)-N-methyltrifluoroacetamide with 1% tert-butylchlorodimethylsilane (Sigma-Aldrich, 00942) and incubation at 60°C for 30 minutes. Gas chromatography coupled to mass spectrometry (GC/MS) was performed using an Agilent 5977C GC/MSD (Agilent, G7077CA) with an Agilent J&W HP-5ms Ultra Inert GC Column (Agilent Technologies, 19091S-433UI). Helium was used as the carrier gas at a constant flow rate of 1.2 mL/min, and one microliter of sample was injected in splitless mode at 250°C. After injection, the GC oven was held at 100°C for 1 min, increased to 105°C at 2.5°C/min, held at 105°C for 2 min, increased to 250°C at 3.5°C/min, and then ramped to 320°C at 20°C/min. The MS system operated under electron impact ionization at 70 eV, and the MS source and quadrupole were held at 230°C and 150°C, respectively. The detector was used in scanning mode with an ion range of 100–650 m/z [133].

#### Intracellular metabolite extraction and analysis via LC/MS

Metabolites were extracted in Eppendorf Safe-Lock tubes (Eppendorf, 022363204) with 80% methanol containing 500 nM of ^13^C/^15^N-labeled Metabolomics Amino Acid Mix (Cambridge Isotope Laboratories, MSK-A2-1.2) as an internal standard. Cells were vortexed vigorously for 10 minutes at 4°C, before pelleting to remove debris and then dried down under nitrogen gas. Metabolites were resuspended in LCMS-grade water, and 2 µL was injected on a Q Exactive^TM^ Benchtop orbitrap mass spectrometer equipped with an Ion Max Source and a HESI II Probe, which was coupled to a Dionex UltiMate3000 HPLC System (ThermoFisher) with a SeQuant® ZIC®-pHILIC 150 x 2.1 mm analytical column equipped with a 2.1 x 20 mm guard column (both 5 mm particle size; Millipore-Sigma). Buffer A was 20 mM ammonium carbonate, 0.1% ammonium hydroxide; Buffer B was acetonitrile. The column oven and autosampler tray were held at 25°C and 4°C, respectively. The chromatographic gradient was run at a flow rate of 0.150 mL/min as follows: 0-20 min: linear gradient from 80-20% B; 20-20.5 min: linear gradient from 20-80% B; 20.5-28 min: hold at 80% B. The mass spectrometer was operated in full-scan, polarity-switching mode, with the spray voltage set to 3.0 kV, the heated capillary held at 275°C, and the HESI probe held at 350°C. The sheath gas flow was set to 40 units, the auxiliary gas flow was set to 15 units, and the sweep gas flow was set to 1 unit. MS data acquisition was performed in a range of *m/z* = 70-1000.

#### Extracellular metabolite quantification from cell culture media by GC-MS

After thawing medium samples on ice, 50 μL of each sample was combined with 10 μL internal standard solution (10 mM U^13^C_6_-D-glucose, 0.1 mM U^13^C_6_-citrate, and 0.5 mM U^13^C_6_,^15^N-L-leucine). Metabolites were extracted with 200 μL of methanol. After vortexing, samples were centrifuged at 14,000 rpm for 10 minutes to remove insoluble material. A volume of 80 μL of the supernatant was transferred to a glass vial and dried under a gentle stream of nitrogen. For lactate quantification, the samples were then incubated with 50 μL of methoxyamine (20 mg/mL in pyridine) at 35°C for 90 minutes, followed by the addition of 50 μL of N-tert-Butyldimethylsilyl-N-methyltrifluoroacetamide (MTBSTFA) at 60°C for 60 minutes. For citrate and pyruvate, the samples were incubated with 50 μL of methoxyamine (20 mg/mL in pyridine) at 35°C for 90 minutes, followed by the addition of 50 μL of N,O-Bis(trimethylsilyl)trifluoroacetamide (BSTFA) at 70°C for 60 minutes. Derivatized samples were analyzed using an Agilent 7890 gas chromatography system coupled to a 5977-mass spectrometer (Agilent Technologies). The injection temperature was set to 280°C with a constant flow rate of 1 mL/minute. Metabolites were separated on a DB-5MS column (Agilent; 30 m x 0.25 mm x 0.25 μM). The oven program for MTBSTFA was set to 80°C, hold for 1 minute, then 15°C/minute to reach 200°C, 10°C/minute to reach 320°C, and hold at 320°C for 6 minutes. For BSTFA analysis, the oven program was set to 60°C, hold for 1 minute, then 10°C/minute to reach 320°C, and hold at 320°C for 5 minutes. Data were acquired in scan mode over a mass range of 50 to 550 Daltons. Data processing was performed using MassHunter Quantitative Analysis software (Agilent Technologies). Metabolites were quantified using external standard calibration curves generated from a series of seven dilutions (0.52 μM to 0.52 mM for citrate, 11 μM to 11 mM for lactate, and 0.11 μM to 0.11 mM for pyruvate). Standards were prepared with the same internal standards as the samples, dried without extraction, and then derivatized and analyzed alongside the experimental samples.

### Quantification and Statistical Analysis

#### Statistical analyses

Statistics were calculated with GraphPad Prism 11 software unless otherwise specified. Unpaired Student’s t-test was used for comparisons between two groups, one-way ANOVA was used for comparisons between >2 groups, and two-way ANOVA was used for grouped comparisons with multiple variables. Graphs display mean values ± standard deviation. All experiments were completed at least twice with three independent biological replicates each time, unless indicated otherwise. The following symbols are used to represent p-values in figures: not significant (ns) for p>0.05, ∗ for p≤0.05, ∗∗ for p≤0.01, ∗∗∗ for p≤0.001, and ∗∗∗∗ for p≤0.0001.

#### Bulk RNA-sequencing analysis

Raw count matrices were obtained from Plasmidsaurus sequencing outputs and processed using R (version 4.4.1) in RStudio (version 2024.4.2.764) with the DESeq2 package (version 1.44.0) for differential expression analysis [134]. Low-abundance genes were excluded by removing features with a total count of ≤5 across all samples. Principal Component Analysis (PCA) was performed on regularized log-transformed (rlog) count data generated using DESeq2 (blind = FALSE), using the PCATools package (version 2.16.0) on the top variable genes (removeVar = 0.1). Transcriptional responses were compared across three carbon source conditions modeled as a categorical variable, and differential expression was assessed using pairwise contrasts with the Wald test. P-values were adjusted for multiple testing using the Benjamini-Hochberg method. Heatmaps were generated from counts per million (CPM) files produced by Plasmidsaurus using the pheatmap package (version 1.0.13). Transcription factor (TF) activity was inferred using the decoupleR package (version 2.10.0) with the univariate linear model (ULM) method [63]. The CollecTRI mouse regulatory network was obtained using the OmniPathR package (version 3.17.7), which was used as prior knowledge to compute TF activity scores with a minimum regulon size of five target genes [135]. Gene Ontology (GO) analysis was performed using ShinyGO 0.85 [136].

#### Metabolomics analysis

Identification and relative quantification of peaks was performed using EL MAVEN software [137]. Identity of all compounds were determined using a library of retention times and MS/MS fragmentation generated from analysis of known standards. For GC-MS, the following ions were utilized for quantification: citrate, 591 *m/z*; α-ketoglutarate, 346 *m/z*; and malate, 419 *m/z*. A mock extraction containing no cells was generated to detect background signal from plastics and sample handling. This mock extraction sample was subtracted from all experimental samples, and peak areas were normalized to internal standards and to live cell count.

#### CCLE bioinformatic Analysis

Gene expression (RNA-seq) and metabolomics data were obtained from the Cancer Cell Line Encyclopedia (CCLE) via the DepMap Public 25Q3 release (Broad Institute). TF activity was inferred as described in “Bulk RNA-sequencing analysis”, except that the Viper algorithm was used. The datasets were then combined, yielding 927 cell lines with both TF activity and metabolomic information. To quantify associations between TF activity and metabolite abundance or derived metabolite ratios, pairwise correlations were assessed using Spearman correlation coefficients (ρ) across all cell lines. Correlation coefficients and corresponding p-values were calculated using cor.test, and p-values were adjusted for multiple hypothesis testing using the Benjamini-Hochberg procedure (p.adjust), which are both functions in base R (stats package). We next assessed whether specific TF programs were significantly positively or negatively enriched for each metabolite or derived metabolite ratio. TFs with padj<0.05 were classified based on the sign of ρ as significantly positively or negatively associated. A pre-defined set of calcium-dependent transcription factors (MEF2A, MEF2B, MEF2C, NFATC2, NFATC1, and CREM) were then tested for enrichment among the positively and negatively associated TFs using Fisher’s exact test by fisher.test (base R stats package). P-values were adjusted by the Benjamini-Hochberg method, and odds ratios were used to quantify the magnitude and direction of enrichment.

#### TCGA Bioinformatic Analysis

Bulk tumor RNA-seq expression data (RSEM values) were downloaded from cBioPortal [138]. The resulting datasets were log2-transformed before calculating pairwise gene-gene Spearman rank correlation coefficients for the following lipid, calcium-response, and T cell lineage genes: SLC25A1, NFATC2, NFATC1, FASN, ACLY, CS, SCD, ACACA, NR4A1, NR4A2, NR4A3, IL2, IL2RA, GZMB, TNF, IFNG, PDCD1, HAVCR2, CD8A, CD8B, FOXP3, and CD4. Correlation coefficients (ρ) and associated two-sided p-values were calculated using the rcorr function in the Hmisc package in R (v. 5.2.5). Gene co-expression networks were constructed in R from pooled TCGA data keeping significant gene pairs meeting thresholds of |ρ| > 0.2 and p<1E-20 as network edges with the igraph (v. 2.2.2) and ggraph (v. 2.2.2) packages. Networks were visualized with node size proportional to degree centrality and edge width proportional to correlation strength. Edge color indicates the sign of the correlation coefficient. Network layouts were generated with a fixed random seed.

#### Spatial Transcriptomics Analysis

Publicly available spatial transcriptomic datasets were curated from DeepSpaceDB [139], only using datasets generated using the 10x Genomics Visium V1 or V2 platforms. Datasets were downloaded and imported into Seurat (version 5.4.0) using the Load10X_Spatial function [140]. Gene expression counts were log normalized in Seurat (NormalizeData). KEGG gene sets were obtained from the MSigDB KEGG Legacy collection. A custom curated gene set for calcium-responsive genes was added for scoring, comprised of: RCAN1, EGR1, EGR2, FOS, FOSB, JUN, JUNB, DUSP1, ATF3, IER2, and BTG2.

Pathway scores were added using the AddModuleScore function in Seurat, keeping pathways with at least 10 genes. Pathway scores were residualized by fitting linear models to each pathway score that included total UMI counts (nCount_Spatial) and gene counts (nFeature_Spatial) as covariates. Spatial cross-correlations between the residualized calcium-response score and each KEGG pathway were evaluated using bivariate Moran’s I statistic by the moran_bv() function in the R package spdep (version 1.4.2). Spatial neighborhoods were defined using a 6-nearest-neighbor graph constructed from tissue coordinates. Empirical two-sided p-values were computed to assess statistical significance using 4,999 random permutations to generate a null distribution of Moran’s I values. P-values were adjusted for multiple hypothesis testing using the Benjamini-Hochberg false discovery rate (FDR). For each KEGG pathway, the frequency of significant spatial co-localization or exclusion with the calcium-response signature (FDR-adjusted p-value < 0.001 and |Moran’s I| > 0.1) was quantified across all datasets.

## Acknowledgements

We thank the Metabolite Profiling Core Facility at the Whitehead Institute for assistance running metabolomics samples, as well as resources provided by the Ragon Institute Flow Cytometry Core. Additionally, we are grateful to Dr. Matthew Vander Heiden and lab members for guidance on method development for gas chromatography-mass spectrometry analyses. We are also grateful to Kendra Libby and Keran Han for feedback and advice. This work was supported by the Koch Institute Support (core) Grant P30-CA14051 from the NCI. L.R.D. was supported by the National Science Foundation Graduate Research Fellowship Program under Grant No. (2141064). Any opinions, findings, and conclusions or recommendations expressed in this material are those of the author(s) and do not necessarily reflect the views of the National Science Foundation. C.L.M. was supported by a fellowship from the Cancer Research Institute. I.J.K and Y.Q. were supported by the Stable Isotope and Metabolomics Core Facility of the Diabetes Research and Training Center (DRTC) of the Albert Einstein College of Medicine (National Institutes of Health, United States; P60DK020541).

## Supplementary Figures and Legends

**Figure S1:**
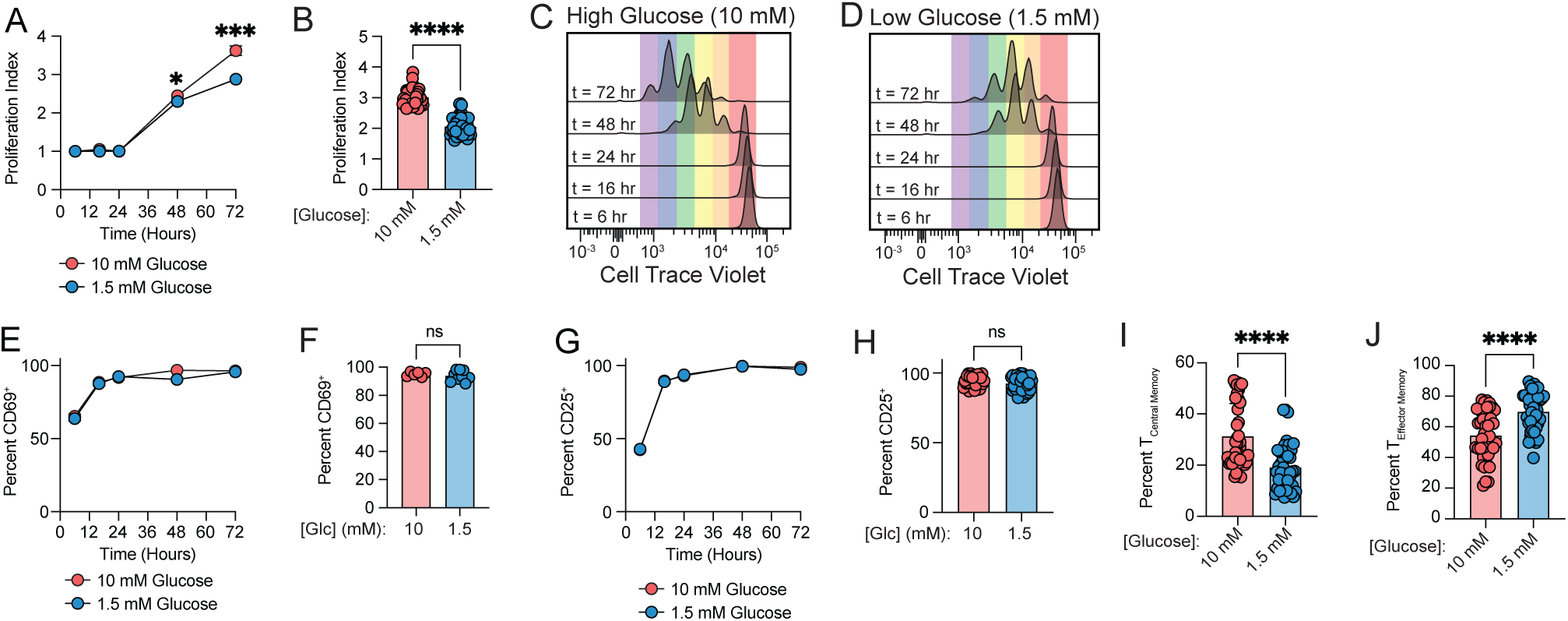
Moderate glucose limitation decreases proliferation and proportion of central memory, but does not affect activation markers. (**A-B**) Proliferation Index value of CD8^+^ T cells at multiple time points during polyclonal stimulation by plate-bound α-CD3/ α-CD28 in high (10 mM) versus low (1.5 mM) glucose (**A**), or after 72 hours (**B**). Proliferation index was calculated using the proliferation modeling tool in FlowJo software. (**C-D**) Representative histograms depicting cell trace violet dye dilution after 72 hours of activation in high glucose (**C**) or low glucose (**D**). (**E-F**) Percent CD69^+^ of live CD8^+^ T cells at multiple time points during polyclonal stimulation by plate-bound α-CD3/ α-CD28 in high (10 mM) versus low (1.5 mM) glucose (**E**), or after 72 hours (**F**). (**G-H**) Percent CD25^+^ of live CD8^+^ T cells at multiple time points during polyclonal stimulation by plate-bound α-CD3/ α-CD28 in high (10 mM) versus low (1.5 mM) glucose (**G**), or after 72 hours (**H**). (**I-J**) Memory state of live CD8^+^ T cells after 72 hours of polyclonal stimulation by plate-bound α-CD3/ α-CD28 in high (10 mM) versus low (1.5 mM) glucose, including central memory (CD62L^+^ CD44^+^) (**I**) and effector memory (CD62L^-^ CD44^+^) (**J**). *Abbreviations*: Glc = glucose; hr = Hours *Statistics*: All data points are distinct biological replicates. Data are presented as mean ± standard deviation (A, B, E-J). Panels A and E contain data representative of two independent experiments where n=3 biological replicates are shown from a representative experiment. Panels B, F, H, I, and J contain data aggregated from 10 independent experiments with n=22 biological replicates. Statistical significance assessed by Student’s t-test (A, B, E-J). P-values: not significant (ns) for p>0.05, ∗ for p≤0.05, ∗∗ for p≤0.01, ∗∗∗ for p≤0.001, and ∗∗∗∗ for p≤0.0001.

**Figure S2:**
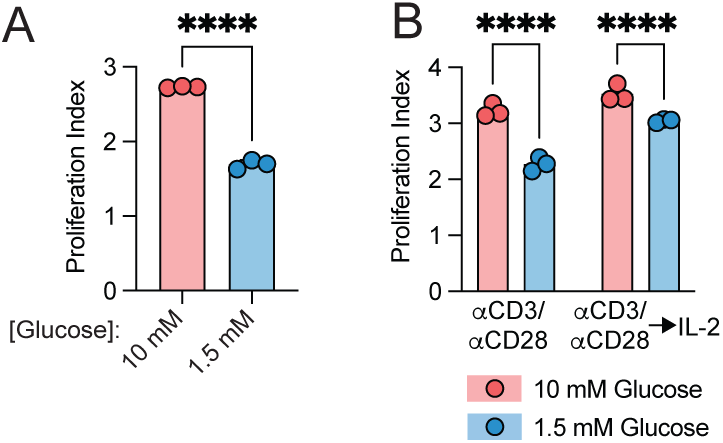
Glucose limitation decreases proliferation across activation modalities of T cell activation. (**A**) Proliferation Index of live CD8^+^ T cells after 72 hours of polyclonal stimulation by plate-bound α-CD3/ α-CD28 in high (10 mM) versus low (1.5 mM) glucose. Proliferation index was calculated using the proliferation modeling tool in FlowJo software. (**B**) Proliferation Index of live CD8^+^ T cells after 24 hours of polyclonal activation on plate-bound α-CD3/ α-CD28 supplemented with IL-2 and then expanded in IL-2 for an additional 72 hours while maintained continuously in high (10 mM) versus low (1.5 mM) glucose. Proliferation index was calculated using the proliferation modeling tool in FlowJo software. *Statistics*: All data points are distinct biological replicates. Data are presented as mean ± standard deviation (A and B). Panels A and B contain data representative of two independent experiments where n=3 biological replicates are shown from a representative experiment. Statistical significance assessed by two-way ANOVA. P-values: not significant (ns) for p>0.05, ∗ for p≤0.05, ∗∗ for p≤0.01, ∗∗∗ for p≤0.001, and ∗∗∗∗ for p≤0.0001.

**Figure S3:**
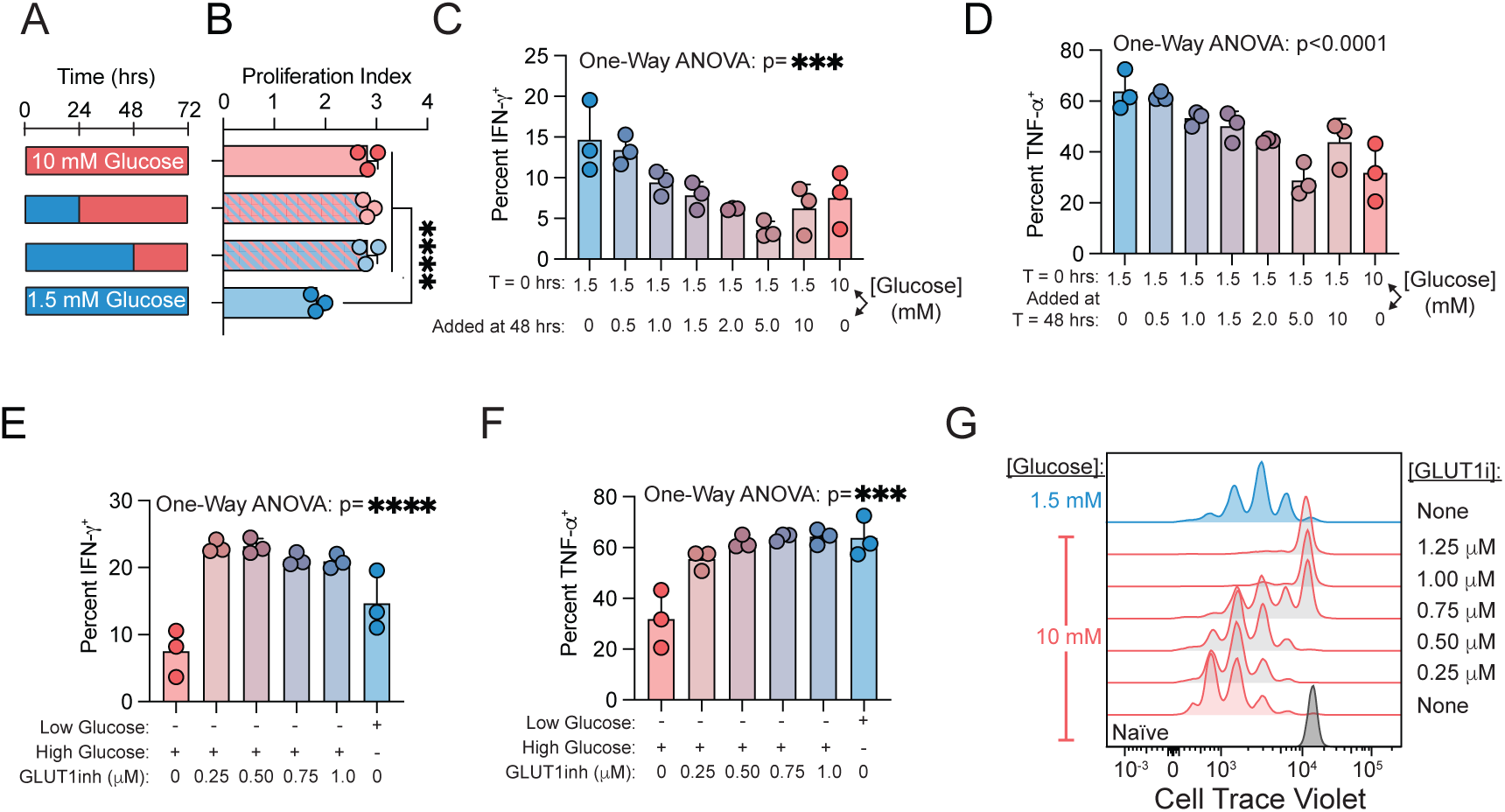
T cell proliferation and production of IFN-γ and TNF-α adapt dynamically to changes in glucose availability. (**A**) Experimental schematic for panel B. (**B**) Proliferation index of live CD8^+^ T cells after 72 hours of activation on plate-bound α-CD3/ α-CD28. Cells were provided glucose at the indicated concentrations and times. Proliferation index was calculated using the proliferation modeling tool in FlowJo software. (C) Percent IFN-γ^+^ of live CD8^+^ T cells activated on plate-bound α-CD3/ α-CD28 in low (1.5 mM) glucose for the first 48 hours before glucose supplementation to the indicated concentrations for the last 24 hours of activation. (D) Percent TNF-α^+^ of live CD8^+^ T cells activated on plate-bound α-CD3/ α-CD28 in low (1.5 mM) glucose for the first 48 hours before glucose supplementation to the indicated concentrations for the last 24 hours of activation. (**E**) Percent IFN-γ^+^ of live CD8^+^ T cells activated on plate-bound α-CD3/ α-CD28 at the indicated glucose concentrations. After 48 hours, a glucose uptake inhibitor (KL-11743) was added for the final 24 hours of activation. (**F**) Percent TNF-α ^+^ of live CD8^+^ T cells activated on plate-bound α-CD3/ α-CD28 at the indicated glucose concentrations. After 48 hours, a glucose uptake inhibitor (KL-11743) was added for the final 24 hours of activation. (**G**) Representative flow cytometry histograms of cell trace violet dye dilution in live CD8^+^ T cells activated on plate-bound α-CD3/ α-CD28 in low (1.5 mM) glucose versus high (10 mM) glucose for the first 48 hours before adding a glucose uptake inhibitor (KL-11743) for the remaining 24 hours of activation at the indicated concentrations. *Abbreviations*: GLUT1inh: Glucose Transporter Inhibitor KL-11743; IFN-γ = Interferon Gamma; TNF-α = Tumor Necrosis Factor alpha. *Statistics*: All data points are distinct biological replicates. Data are presented as mean ± standard deviation (B-F). Panels B-F contain data representative of two independent experiments where n=3 biological replicates are shown from a representative experiment. Statistical significance assessed by one-way ANOVA (B) or two-way ANOVA (C-F). P-values: not significant (ns) for p>0.05, ∗ for p≤0.05, ∗∗ for p≤0.01, ∗∗∗ for p≤0.001, and ∗∗∗∗ for p≤0.0001.

**Figure S4:**
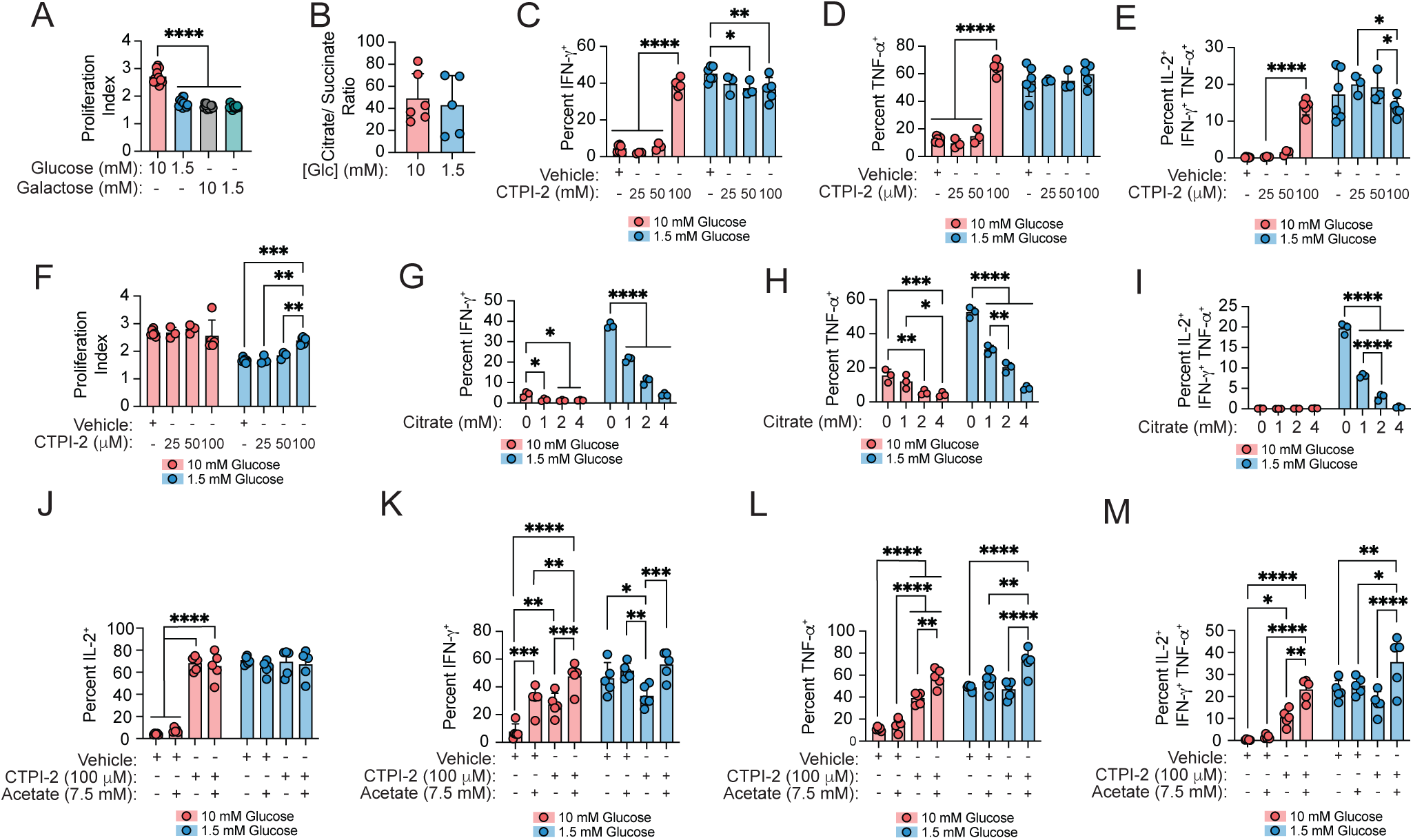
IFN-γ, TNF-α, and cell proliferation levels are limited by the export of citrate from the mitochondria via SLC25A1. (**A**) Proliferation Index of live CD8^+^ T cells after 72 hours of polyclonal activation by plate-bound α-CD3/ α-CD28 in glucose or galactose at the indicated concentrations. Proliferation index was calculated using the proliferation modeling tool in FlowJo software. (**B**) Metabolomics analysis on CD8^+^ T cells after 72 hours of activation by plate-bound α-CD3/ α-CD28 in high (10 mM) versus low (1.5 mM) glucose measured by liquid chromatography-mass spectrometry (LC-MS). Barplots depict ratios of peak areas for citrate to succinate. (**C-E**) Percent IFN-γ^+^ (**C**), TNF-α^+^ (**D**), and polyfunctional (IL-2^+^ IFN-γ^+^ TNF-α^+^) (**E**) of live CD8^+^ T cells after 72 hours of polyclonal activation by plate-bound α-CD3/ α-CD28 in high (10 mM) versus low (1.5 mM) glucose with varying concentrations of the mitochondrial citrate carrier inhibitor (SLC25A1) inhibitor CTPI-2 or vehicle control. (**F**) Proliferation Index of live CD8^+^ T cells after 72 hours of polyclonal activation by plate-bound α-CD3/ α-CD28 in high (10 mM) versus low (1.5 mM) glucose with varying concentrations of the mitochondrial citrate carrier inhibitor (SLC25A1) inhibitor CTPI-2 or vehicle control. (**G-I**) Percent IFN-γ^+^ (**G**), TNF-α^+^ (**H**), and polyfunctional (IL-2^+^ IFN-γ^+^ TNF-α^+^) (**I**) of live CD8^+^ T cells after 72 hours of polyclonal activation by plate-bound α-CD3/ α-CD28 in high (10 mM) versus low (1.5 mM) glucose with varying concentrations of sodium citrate. (**J-M**) Percent IL-2^+^ (**J**), IFN-γ^+^ (**K**), TNF-α^+^ (**L**), and polyfunctional (IL-2^+^ IFN-γ^+^ TNF-α^+^) (**M**) of live CD8^+^ T cells after 72 hours of polyclonal activation by plate-bound α-CD3/ α-CD28 in high (10 mM) versus low (1.5 mM) glucose, with addition of 7.5 mM sodium acetate alone or in combination with 100 μM of the mitochondrial citrate carrier inhibitor (SLC25A1) inhibitor CTPI-2. *Abbreviations*: Glc = Glucose; IFN-γ = Interferon Gamma; TNF-α = Tumor Necrosis Factor alpha; CTPI-2 = Inhibitor of the Mitochondrial Citrate Carrier SLC25A1. *Statistics*: Data are presented as mean ± standard deviation (A-M). Panel A contains data aggregated from 5 independent experiments with n=15 biological replicates. Panel B contains data aggregated from 2 independent experiments with n=6 biological replicates with 5 technical replicates each. Panels C-F contain data aggregated from 3 independent experiments with n=7 biological replicates. Panels G-I contain data representative of two independent experiments where n=3 biological replicates are shown. Panels J-M contain data representative of two independent experiments where n=5 biological replicates are shown. Statistical significance assessed by one-way ANOVA (A), Student’s t-test (B), or two-way ANOVA (C-M). P-values: not significant (ns) for p>0.05, ∗ for p≤0.05, ∗∗ for p≤0.01, ∗∗∗ for p≤0.001, and ∗∗∗∗ for p≤0.0001.

**Figure S5:**
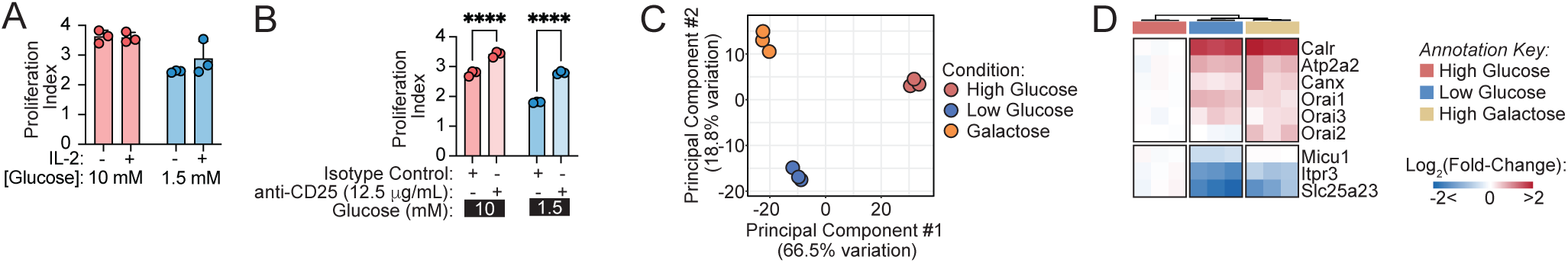
Cellular responses to glucose restriction or replacement with galactose. (**A**) Proliferation index of live CD8^+^ T cells after 72 hours of polyclonal activation on plate-bound α-CD3/ α-CD28 in high (10 mM) versus low (1.5 mM) glucose. Exogenous IL-2 is supplemented as indicated. Proliferation index was calculated using the proliferation modeling tool in FlowJo software. (**B**) Proliferation index of live CD8^+^ T cells after 72 hours of polyclonal activation on plate-bound α-CD3/ CD28 in high (10 mM) versus low (1.5 mM) glucose. A CD25-blocking antibody or isotype control are added as indicated. Proliferation index was calculated using the proliferation modeling tool in FlowJo software. (C) Principal component analysis plot of differential gene analysis from bulk RNA sequencing of CD8^+^ T cells after 72 hours of activation in high (10 mM) glucose, low (1.5 mM) glucose, or galactose. (D) Heatmap highlighting differentially expressed genes from bulk RNA sequencing of CD8^+^ T cells that participate in calcium intake and handling. *Statistics*: All data points are distinct biological replicates. Data are presented as mean ± standard deviation (A-B). Panels A-B contain data representative of two independent experiments where n=3 biological replicates are shown from a representative experiment. Statistical significance assessed by two-way ANOVA (A-B). P-values: not significant (ns) for p>0.05, ∗ for p≤0.05, ∗∗ for p≤0.01, ∗∗∗ for p≤0.001, and ∗∗∗∗ for p≤0.0001.

**Figure S6:**
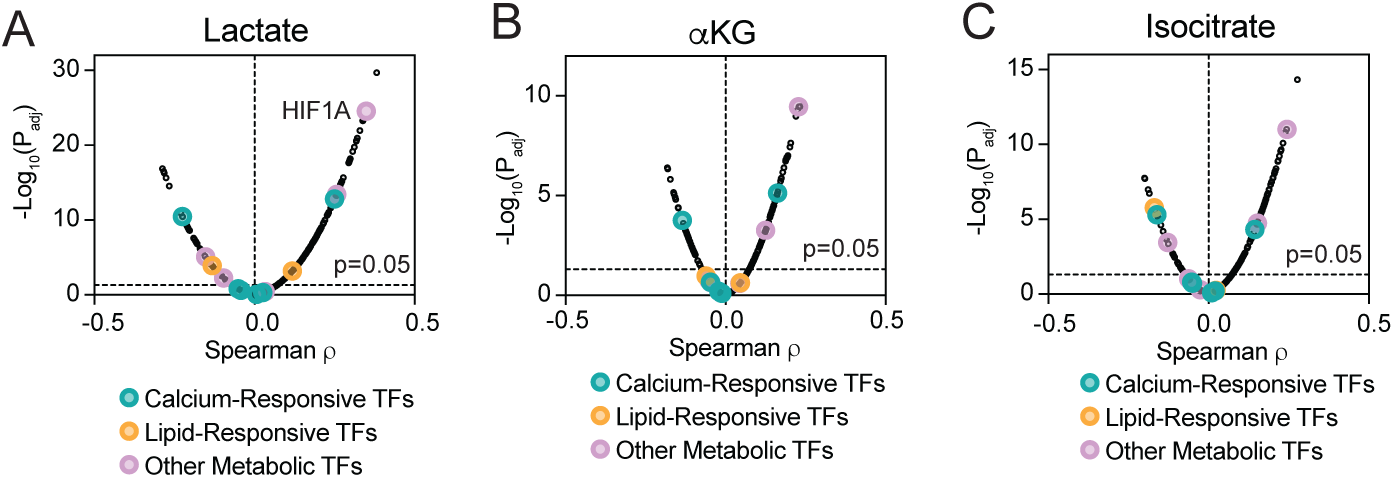
Citrate, but not other metabolites generated during mitochondrial glucose oxidation, is negatively associated with calcium-dependent transcriptional programs. (**A-C**) Positive and negative associations between transcription factor programs and metabolite levels, including lactate (**A**), α-Ketoglutarate (**B**), and isocitrate (**C**) within cancer cell lines from the Cancer Cell Line Encyclopedia (CCLE). *Abbreviations*: HIF1A = Hypoxia-Inducible Factor 1-Alpha; CCLE = Cancer Cell Line Encyclopedia; TFs = Transcription Factors.

## References

1. Wang, R., Dillon, C.P., Shi, L.Z., Milasta, S., Carter, R., Finkelstein, D., McCormick, L.L., Fitzgerald, P., Chi, H., Munger, J., et al. (2011). The transcription factor Myc controls metabolic reprogramming upon T lymphocyte activation. Immunity 35, 871. 10.1016/J.IMMUNI.2011.09.021.

2. Frauwirth, K.A., Riley, J.L., Harris, M.H., Parry, R. V., Rathmell, J.C., Plas, D.R., Elstrom, R.L., June, C.H., and Thompson, C.B. (2002). The CD28 signaling pathway regulates glucose metabolism. Immunity 16, 769–777. 10.1016/S1074-7613(02)00323-0/ASSET/C91BE218-FAB7-4281-A125-C7C2D8B0681B/MAIN.ASSETS/GR7.JPG.

3. Jacobs, S.R., Herman, C.E., MacIver, N.J., Wofford, J.A., Wieman, H.L., Hammen, J.J., and Rathmell, J.C. (2008). Glucose Uptake Is Limiting in T Cell Activation and Requires CD28-Mediated Akt-Dependent and Independent Pathways. J. Immunol. 180, 4476. 10.4049/JIMMUNOL.180.7.4476.

4. Jones, R.G., and Thompson, C.B. (2007). Revving the Engine: Signal Transduction Fuels T Cell Activation. Immunity 27, 173–178. 10.1016/j.immuni.2007.07.008.

5. D’Souza, A.D., Parikh, N., Kaech, S.M., and Shadel, G.S. (2007). Convergence of multiple signaling pathways is required to coordinately up-regulate mtDNA and mitochondrial biogenesis during T cell activation. Mitochondrion 7, 374. 10.1016/J.MITO.2007.08.001.

6. Ron-Harel, N., Santos, D., Ghergurovich, J.M., Sage, P.T., Reddy, A., Lovitch, S.B., Dephoure, N., Satterstrom, F.K., Sheffer, M., Spinelli, J.B., et al. (2016). Mitochondrial Biogenesis and Proteome Remodeling Promote One-Carbon Metabolism for T Cell Activation. Cell Metab. 24, 104–117. 10.1016/j.cmet.2016.06.007.

7. Fox, C.J., Hammerman, P.S., and Thompson, C.B. (2005). Fuel feeds function: energy metabolism and the T-cell response. Nature Reviews Immunology 2005 5:11 5, 844–852. 10.1038/NRI1710.

8. Gerriets, V.A., and Rathmell, J.C. (2012). Metabolic Pathways in T Cell Fate and Function. Trends Immunol. 33, 168. 10.1016/J.IT.2012.01.010.

9. Heiden, M.G.V., Cantley, L.C., and Thompson, C.B. (2009). Understanding the Warburg effect: the metabolic requirements of cell proliferation. Science 324, 1029–1033. 10.1126/SCIENCE.1160809.

10. Warburg, O. (1956). On the origin of cancer cells. Science 123, 309–314. 10.1126/SCIENCE.123.3191.309.

11. Raynor, J.L., and Chi, H. (2024). Nutrients: Signal 4 in T cell immunity. J. Exp. Med. 221, e20221839. 10.1084/JEM.20221839.

12. Roos, D., and Loos, J.A. (1970). Changes in the carbohydrate metabolism of mitogenicellay stimulated human peripheral lymphocytes I. Stimulation by phytohaemagglutinin. Biochimica et Biophysica Acta (BBA) - General Subjects 222, 565–582. 10.1016/0304-4165(70)90182-0.

13. Hedeskov, C.J. (1968). Early effects of phytohaemagglutinin on glucose metabolism of normal human lymphocytes. Biochemical Journal 110, 373. 10.1042/BJ1100373.

14. Culvenor, J.G., and Weidemann, M.J. (1976). Phytohaemagglutinin stimulation of rat thymus lymphocyte glycolysis. Biochimica et Biophysica Acta (BBA) - General Subjects 437, 354–363. 10.1016/0304-4165(76)90005-2.

15. Cooper, E.H., Barkhan, P., and Hale, A.J. (1963). Observations on the Proliferation of Human Leucocytes Cultured with Phytohaemagglutinin. Br. J. Haematol. 9, 101–111. 10.1111/J.1365-2141.1963.TB05446.X.

16. Menk, A. V., Scharping, N.E., Moreci, R.S., Zeng, X., Guy, C., Salvatore, S., Bae, H., Xie, J., Young, H.A., Wendell, S.G., et al. (2018). Early TCR Signaling Induces Rapid Aerobic Glycolysis Enabling Distinct Acute T Cell Effector Functions. Cell Rep. 22, 1509. 10.1016/J.CELREP.2018.01.040.

17. Ma, S., Ming, Y., Wu, J., and Cui, G. (2024). Cellular metabolism regulates the differentiation and function of T-cell subsets. Cellular & Molecular Immunology 2024 21:5 21, 419–435. 10.1038/s41423-024-01148-8.

18. Michalek, R.D., Gerriets, V.A., Jacobs, S.R., Macintyre, A.N., MacIver, N.J., Mason, E.F., Sullivan, S.A., Nichols, A.G., and Rathmell, J.C. (2011). Cutting edge: distinct glycolytic and lipid oxidative metabolic programs are essential for effector and regulatory CD4+ T cell subsets. J. Immunol. 186, 3299–3303. 10.4049/JIMMUNOL.1003613.

19. Shi, L.Z., Wang, R., Huang, G., Vogel, P., Neale, G., Green, D.R., and Chi, H. (2011). HIF1α–dependent glycolytic pathway orchestrates a metabolic checkpoint for the differentiation of TH17 and Treg cells. J. Exp. Med. 208, 1367. 10.1084/JEM.20110278.

20. Xu, K., Yin, N., Peng, M., Stamatiades, E.G., Chhangawala, S., Shyu, A., Li, P., Zhang, X., Do, M.H., Capistrano, K.J., et al. (2021). Glycolytic ATP Fuels Phosphoinositide 3-Kinase Signaling to Support Effector T Helper 17 Cell Responses. Immunity 54, 976. 10.1016/J.IMMUNI.2021.04.008.

21. Xu, K., Yin, N., Peng, M., Stamatiades, E.G., Shyu, A., Li, P., Zhang, X., Do, M.H., Wang, Z., Capistrano, K.J., et al. (2021). Glycolysis fuels phosphoinositide 3-kinase signaling to bolster T cell immunity. Science 371, 405–410. 10.1126/SCIENCE.ABB2683.

22. Ho, P.C., Bihuniak, J.D., MacIntyre, A.N., Staron, M., Liu, X., Amezquita, R., Tsui, Y.C., Cui, G., Micevic, G., Perales, J.C., et al. (2015). Phosphoenolpyruvate Is a Metabolic Checkpoint of Anti-tumor T Cell Responses. Cell 162, 1217–1228. 10.1016/J.CELL.2015.08.012.

23. Su, E.W., Moore, C.J., Suriano, S., Johnson, C.B., Songalia, N., Patterson, A., Neitzke, D.J., Andrijauskaite, K., Garrett-Mayer, E., Mehrotra, S., et al. (2015). IL-2Rα mediates temporal regulation of IL-2 signaling and enhances immunotherapy. Sci. Transl. Med. 7, 311ra170. 10.1126/SCITRANSLMED.AAC8155.

24. Cham, C.M., Driessens, G., O’Keefe, J.P., and Gajewski, T.F. (2008). Glucose deprivation inhibits multiple key gene expression events and effector functions in CD8+ T cells. Eur. J. Immunol. 38, 2438–2450. 10.1002/EJI.200838289.

25. Cham, C.M., and Gajewski, T.F. (2005). Glucose Availability Regulates IFN-γ Production and p70S6 Kinase Activation in CD8+ Effector T Cells. The Journal of Immunology 174, 4670–4677. 10.4049/JIMMUNOL.174.8.4670.

26. Chang, C.H., Curtis, J.D., Maggi, L.B., Faubert, B., Villarino, A. V., O’Sullivan, D., Huang, S.C.C., Van Der Windt, G.J.W., Blagih, J., Qiu, J., et al. (2013). Posttranscriptional Control of T Cell Effector Function by Aerobic Glycolysis. Cell 153, 1239. 10.1016/J.CELL.2013.05.016.

27. Sena, L.A., Li, S., Jairaman, A., Prakriya, M., Ezponda, T., Hildeman, D.A., Wang, C.R., Schumacker, P.T., Licht, J.D., Perlman, H., et al. (2013). Mitochondria are required for antigen-specific T cell activation through reactive oxygen species signaling. Immunity 38, 225. 10.1016/J.IMMUNI.2012.10.020.

28. Blagih, J., Coulombe, F., Vincent, E.E., Dupuy, F., Galicia-Vázquez, G., Yurchenko, E., Raissi, T.C., vanderWindt, G.J.W., Viollet, B., Pearce, E.L., et al. (2015). The energy sensor AMPK regulates T cell metabolic adaptation and effector responses in vivo. Immunity 42, 41–54. 10.1016/J.IMMUNI.2014.12.030.

29. Jacobs, S.R., Herman, C.E., Maciver, N.J., Wofford, J.A., Wieman, H.L., Hammen, J.J., and Rathmell, J.C. (2008). Glucose Uptake Is Limiting in T Cell Activation and Requires CD28-Mediated Akt-Dependent and Independent Pathways 1.

30. Chang, C.H., Curtis, J.D., Maggi, L.B., Faubert, B., Villarino, A. V., O’Sullivan, D., Huang, S.C.C., Van Der Windt, G.J.W., Blagih, J., Qiu, J., et al. (2013). XPosttranscriptional control of T cell effector function by aerobic glycolysis. Cell 153, 1239. 10.1016/j.cell.2013.05.016.

31. Frisch, A.T., Wang, Y., Xie, B., Yang, A., Ford, B.R., Joshi, S., Kedziora, K.M., Peralta, R., Wilfahrt, D., Mullett, S.J., et al. (2025). Redirecting glucose flux during in vitro expansion generates epigenetically and metabolically superior T cells for cancer immunotherapy. Cell Metab. 37, 870. 10.1016/J.CMET.2024.12.007.

32. Li, H., Ning, S., Ghandi, M., Kryukov, G. V., Gopal, S., Deik, A., Souza, A., Pierce, K., Keskula, P., Hernandez, D., et al. (2019). The Landscape of Cancer Cell Line Metabolism. Nat. Med. 25, 850. 10.1038/S41591-019-0404-8.

33. Greiners, E.F., Guppy9, M., and Brandh, K. (1994). T H E JOURNAL OF BIOLOGICAL CHEMISTRY Glucose Is Essential for Proliferation and the Glycolytic Enzyme Induction That Provokes a Transition to Glycolytic Energy Production*. Journal of Biological Chemistry 269, 31484–31490. 10.1016/S0021-9258(18)31720-4.

34. Qin, M., Shang, L., Chen, H., Shi, L., Liu, C., Ding, M., He, D., Shao, C., Yuan, S., Yu, H., et al. (2026). SLC13A2-transported citrate remodels transcriptional regulation through protein acetylation to suppress tumor growth. Sci. Adv. 12, eaec4368. 10.1126/SCIADV.AEC4368/SUPPL_FILE/SCIADV.AEC4368_SM.PDF.

35. Fu, H., Vuononvirta, J., Fanti, S., Bonacina, F., D’Amati, A., Wang, G., Poobalasingam, T., Fankhaenel, M., Lucchesi, D., Coleby, R., et al. (2023). The glucose transporter 2 regulates CD8+ T cell function via environment sensing. Nature Metabolism 2023 5:11 5, 1969–1985. 10.1038/s42255-023-00913-9.

36. Wals, P.A., and Katz, J. (1993). A concentration gradient of glucose from liver to plasma. Metabolism 42, 1492–1496. 10.1016/0026-0495(93)90204-2.

37. Goodwin, M.L. (2010). Blood Glucose Regulation during Prolonged, Submaximal, Continuous Exercise: A Guide for Clinicians. J. Diabetes Sci. Technol. 4, 694. 10.1177/193229681000400325.

38. 2. Diagnosis and Classification of Diabetes: Standards of Care in Diabetes-2026 (2026). Diabetes Care 49, S27–S49. 10.2337/DC26-S002.

39. Regittnig, W., Ellmerer, M., Fauler, G., Sendlhofer, G., Trajanoski, Z., Leis, H.J., Schaupp, L., Wach, P., and Pieber, T.R. (2003). Assessment of transcapillary glucose exchange in human skeletal muscle and adipose tissue. Am. J. Physiol. Endocrinol. Metab. 285. 10.1152/AJPENDO.00351.2002.

40. MacLean, D.A., Bangsbo, J., and Saltin, B. (1999). Muscle interstitial glucose and lactate levels during dynamic exercise in humans determined by microdialysis. J. Appl. Physiol. (1985) 87, 1483–1490. 10.1152/JAPPL.1999.87.4.1483.

41. Maggs, D.G., Jacob, R., Rife, F., Lange, R., Leone, P., During, M.J., Tamborlane, W. V., and Sherwin, R.S. (1995). Interstitial fluid concentrations of glycerol, glucose, and amino acids in human quadricep muscle and adipose tissue. Evidence for significant lipolysis in skeletal muscle. J. Clin. Invest. 96, 370–377. 10.1172/JCI118043.

42. Apiz Saab, J.J., and Muir, A. (2023). Tumor interstitial fluid analysis enables the study of microenvironment-cell interactions in cancers. Curr. Opin. Biotechnol. 83. 10.1016/J.COPBIO.2023.102970.

43. Wherry, E.J., Blattman, J.N., Murali-Krishna, K., van der Most, R., and Ahmed, R. (2003). Viral Persistence Alters CD8 T-Cell Immunodominance and Tissue Distribution and Results in Distinct Stages of Functional Impairment. J. Virol. 77, 4911. 10.1128/JVI.77.8.4911-4927.2003.

44. D’Souza, W.N., and Lefrançois, L. (2004). Frontline: An in-depth evaluation of the production of IL-2 by antigen-specific CD8 T cells in vivo. Eur. J. Immunol. 34, 2977–2985. 10.1002/EJI.200425485.

45. Hogquist, K.A., Jameson, S.C., Heath, W.R., Howard, J.L., Bevan, M.J., and Carbone, F.R. (1994). T cell receptor antagonist peptides induce positive selection. Cell 76, 17–27. 10.1016/0092-8674(94)90169-4.

46. Caputto, R., and Leloir, L.R. (1949). THE ENZYMATIC TRANSFORMATION OF GALACTOSE INTO GLUCOSE DERIVATIVES. Journal of Biological Chemistry 179, 497–498. 10.1016/S0021-9258(18)56863-0.

47. Rossignol, R., Gilkerson, R., Aggeler, R., Yamagata, K., Remington, S.J., and Capaldi, R.A. (2004). Energy substrate modulates mitochondrial structure and oxidative capacity in cancer cells. Cancer Res. 64, 985–993. 10.1158/0008-5472.CAN-03-1101.

48. Bustamente, E., Morris, H.P., and Pedersen, P.L. (1977). Hexokinase: the direct link between mitochondrial and glycolytic reactions in rapidly growing cancer cells. Adv. Exp. Med. Biol. 92, 363–380. 10.1007/978-1-4615-8852-8_15.

49. Weinberg, F., Hamanaka, R., Wheaton, W.W., Weinberg, S., Joseph, J., Lopez, M., Kalyanaraman, B., Mutlu, G.M., Budinger, G.R.S., and Chandel, N.S. (2010). Mitochondrial metabolism and ROS generation are essential for Kras-mediated tumorigenicity. Proc. Natl. Acad. Sci. U. S. A. 107, 8788–8793. 10.1073/PNAS.1003428107/-/DCSUPPLEMENTAL.

50. Kaymak, I., Watson, M.J., Oswald, B.M., Ma, S., Johnson, B.K., Decamp, L.M., Mabvakure, B.M., Luda, K.M., Ma, E.H., Lau, K., et al. (2024). ACLY and ACSS2 link nutrient-dependent chromatin accessibility to CD8 T cell effector responses. Journal of Experimental Medicine 221. 10.1084/JEM.20231820/276919.

51. Wellen, K.E., Hatzivassiliou, G., Sachdeva, U.M., Bui, T. V., Cross, J.R., and Thompson, C.B. (2009). ATP-citrate lyase links cellular metabolism to histone acetylation. Science 324, 1076–1080. 10.1126/SCIENCE.1164097.

52. Berwick, D.C., Hers, I., Heesom, K.J., Kelly Moule, S., and Tavaré, J.M. (2002). The identification of ATP-citrate lyase as a protein kinase B (Akt) substrate in primary adipocytes. J. Biol. Chem. 277, 33895–33900. 10.1074/JBC.M204681200.

53. Arnold, P.K., Jackson, B.T., Paras, K.I., Brunner, J.S., Hart, M.L., Newsom, O.J., Alibeckoff, S.P., Endress, J., Drill, E., Sullivan, L.B., et al. (2022). A non-canonical tricarboxylic acid cycle underlies cellular identity. Nature 2022 603:7901 603, 477–481. 10.1038/s41586-022-04475-w.

54. Xie, A., Brunner, J.S., Chakraborty, S., Montero, A.M., Bridgeman, A.E., Paras, K.I., Cui, R., Fagoaga-Eugui, M., Komza, M., Arnold, P.K., et al. (2026). Citrate clearance is a major function of aconitase 2 in the canonical TCA cycle. Cell 189, 2684. 10.1016/J.CELL.2026.01.028.

55. Wieland, O.H. (1983). The mammalian pyruvate dehydrogenase complex: structure and regulation. Rev. Physiol. Biochem. Pharmacol. 96, 123–170. 10.1007/BFB0031008/SAVE-RESEARCH.

56. Stipani, I., Krämer, R., Palmieri, F., and Klingenberg, M. (1980). Citrate transport in liposomes reconstituted with triton extracts from mitochondria. Biochem. Biophys. Res. Commun. 97, 1206–1214. 10.1016/0006-291X(80)91503-X.

57. Stipani, I., and Palmieri, F. (1983). Purification of the active mitochondrial tricarboxylate carrier by hydroxylapatite chromatography. FEBS Lett. 161, 269–274. 10.1016/0014-5793(83)81023-0.

58. Bisaccia, F., De Palma, A., and Palmieri, F. (1989). Identification and purification of the tricarboxylate carrier from rat liver mitochondria. Biochim. Biophys. Acta 977, 171–176. 10.1016/S0005-2728(89)80068-4.

59. Kahan, S.M., Bakshi, R.K., Ingram, J.T., Hendrickson, R.C., Lefkowitz, E.J., Crossman, D.K., Harrington, L.E., Weaver, C.T., and Zajac, A.J. (2022). Intrinsic IL-2 Production by Effector CD8 T Cells Affects IL-2 Signaling and Promotes Fate Decisions, Stemness, and Protection. Sci. Immunol. 7, eabl6322. 10.1126/SCIIMMUNOL.ABL6322.

60. Halban, P.A., and Irminger, J.C. (1994). Sorting and processing of secretory proteins. Biochemical Journal 299, 1. 10.1042/BJ2990001.

61. McGraw, C.F., Somlyo, A. V., and Blaustein, M.P. (1980). Localization of calcium in presynaptic nerve terminals. An ultrastructural and electron microprobe analysis. J. Cell Biol. 85, 228–241. 10.1083/JCB.85.2.228.

62. Berridge, M.J. (1993). Inositol trisphosphate and calcium signalling. Nature 361, 315–325. 10.1038/361315A0.

63. Badia-I-Mompel, P., Vélez Santiago, J., Braunger, J., Geiss, C., Dimitrov, D., Müller-Dott, S., Taus, P., Dugourd, A., Holland, C.H., Ramirez Flores, R.O., et al. (2022). decoupleR: ensemble of computational methods to infer biological activities from omics data. Bioinformatics advances 2. 10.1093/BIOADV/VBAC016.

64. Shaw, J.P., Utz, P.J., Durand, D.B., Toole, J.J., Emmel, E.A., and Crabtree, G.R. (1988). Identification of a Putative Regulator of Early T Cell Activation Genes. Science (1979). 185, 4972–4975. 10.1126/SCIENCE.3260404.

65. Chen, L., Glovert, J.N.M., Hogan, P.G., Rao, A., and Harrison, S.C. (1998). Structure of the DNA-binding domains from NFAT, Fos and Jun bound specifically to DNA. Nature 1998 392:6671 392, 42–48. 10.1038/32100.

66. Okamura, H., Aramburu, J., García-Rodríguez, C., Viola, J.P.B., Raghavan, A., Tahiliani, M., Zhang, X., Qin, J., Hogan, P.G., and Rao, A. (2000). Concerted dephosphorylation of the transcription factor NFAT1 induces a conformational switch that regulates transcriptional activity. Mol. Cell 6, 539–550. 10.1016/S1097-2765(00)00053-8.

67. Zomorodian, A., and Moe, O.W. (2025). Citrate and calcium kidney stones. Clin. Kidney J. 18, sfaf244. 10.1093/CKJ/SFAF244.

68. Hu, Y.Y., Rawal, A., and Schmidt-Rohr, K. (2010). Strongly bound citrate stabilizes the apatite nanocrystals in bone. Proc. Natl. Acad. Sci. U. S. A. 107, 22425–22429. 10.1073/PNAS.1009219107/-/DCSUPPLEMENTAL.

69. Singh, K.K., Desouki, M.M., Franklin, R.B., and Costello, L.C. (2006). Mitochondrial aconitase and citrate metabolism in malignant and nonmalignant human prostate tissues. Mol. Cancer 5, 14-. 10.1186/1476-4598-5-14/FIGURES/3.

70. Xie, A., Brunner, J.S., Chakraborty, S., Montero, A.M., Bridgeman, A.E., Paras, K.I., Cui, R., Fagoaga-Eugui, M., Komza, M., Arnold, P.K., et al. (2026). Citrate clearance is a major function of aconitase 2 in the canonical TCA cycle. Cell 189, 2684. 10.1016/J.CELL.2026.01.028.

71. Barretina, J., Caponigro, G., Stransky, N., Venkatesan, K., Margolin, A.A., Kim, S., Wilson, C.J., Lehár, J., Kryukov, G. V., Sonkin, D., et al. (2012). The Cancer Cell Line Encyclopedia enables predictive modelling of anticancer drug sensitivity. Nature 483, 603–607. 10.1038/NATURE11003.

72. Semenza, G.L., Jiang, B.H., Leung, S.W., Passantino, R., Concordat, J.P., Maire, P., and Giallongo, A. (1996). Hypoxia response elements in the aldolase A, enolase 1, and lactate dehydrogenase A gene promoters contain essential binding sites for hypoxia-inducible factor 1. J. Biol. Chem. 271, 32529–32537. 10.1074/JBC.271.51.32529.

73. Seagroves, T.N., Ryan, H.E., Lu, H., Wouters, B.G., Knapp, M., Thibault, P., Laderoute, K., and Johnson, R.S. (2001). Transcription factor HIF-1 is a necessary mediator of the pasteur effect in mammalian cells. Mol. Cell. Biol. 21, 3436–3444. 10.1128/MCB.21.10.3436-3444.2001.

74. Iyer, N. V., Kotch, L.E., Agani, F., Leung, S.W., Laughner, E., Wenger, R.H., Gassmann, M., Gearhart, J.D., Lawler, A.M., Yu, A.Y., et al. (1998). Cellular and developmental control of O2 homeostasis by hypoxia-inducible factor 1 alpha. Genes Dev. 12, 149–162. 10.1101/GAD.12.2.149.

75. Chypre, M., Zaidi, N., and Smans, K. (2012). ATP-citrate lyase: A mini-review. Biochem. Biophys. Res. Commun. 422, 1–4. 10.1016/J.BBRC.2012.04.144.

76. Srere, P.A. (1959). The Citrate Cleavage Enzyme: I. DISTRIBUTION AND PURIFICATION. Journal of Biological Chemistry 234, 2544–2547. 10.1016/S0021-9258(18)69735-2.

77. Tan, M., Mosaoa, R., Graham, G.T., Kasprzyk-Pawelec, A., Gadre, S., Parasido, E., Catalina-Rodriguez, O., Foley, P., Giaccone, G., Cheema, A., et al. (2020). Inhibition of the mitochondrial citrate carrier, Slc25a1, reverts steatosis, glucose intolerance, and inflammation in preclinical models of NAFLD/NASH. Cell Death & Differentiation 2020 27:7 27, 2143–2157. 10.1038/s41418-020-0491-6.

78. Yang, Y., He, J., Zhang, B., Zhang, Z., Jia, G., Liu, S., Wu, T., He, X., and Wang, N. (2021). SLC25A1 promotes tumor growth and survival by reprogramming energy metabolism in colorectal cancer. Cell Death & Disease 2021 12:12 12, 1108-. 10.1038/s41419-021-04411-2.

79. Weinstein, J.N., Collisson, E.A., Mills, G.B., Shaw, K.R.M., Ozenberger, B.A., Ellrott, K., Sander, C., Stuart, J.M., Chang, K., Creighton, C.J., et al. (2013). The Cancer Genome Atlas Pan-Cancer analysis project. Nature Genetics 2013 45:10 45, 1113–1120. 10.1038/ng.2764.

80. Kanehisa, M., Furumichi, M., Tanabe, M., Sato, Y., and Morishima, K. (2017). KEGG: new perspectives on genomes, pathways, diseases and drugs. Nucleic Acids Res. 45, D353–D361. 10.1093/NAR/GKW1092.

81. Whitfield, J.F., Perris, A.D., and Youdale, T. (1968). The role of calcium in the mitotic stimulation of rat thymocytes by detergents, agmatine and poly-l-lysine. Exp. Cell Res. 53, 155–165. 10.1016/0014-4827(68)90363-7.

82. Asherson, G.L., Davey, M.J., and Goodford, P.J. (1970). Increased uptake of calcium by human lymphocytes treated with phytohaemagglutinin. J. Physiol. 206, 32P–33P.

83. Trebak, M., and Kinet, J.P. (2019). Calcium signalling in T cells. Nature Reviews Immunology 2018 19:3 19, 154–169. 10.1038/S41577-018-0110-7.

84. Gouy, H., Cefai, D., Christensen, S.B., Debré, P., and Bismuth, G. (1990). Ca2+ influx in human T lymphocytes is induced independently of inositol phosphate production by mobilization of intracellular Ca2+ stores. A study with the Ca2+ endoplasmic reticulum-ATPase inhibitor thapsigargin. Eur. J. Immunol. 20, 2269–2275. 10.1002/EJI.1830201016.

85. Feske, S., Giltnane, J., Dolmetsch, R., Staudt, L.M., and Rao, A. (2001). Gene regulation mediated by calcium signals in T lymphocytes. Nature Immunology 2001 2:4 2, 316–324. 10.1038/86318.

86. Macian, F. (2005). NFAT proteins: key regulators of T-cell development and function. Nature Reviews Immunology 2005 5:6 5, 472–484. 10.1038/nri1632.

87. Feske, S., Skolnik, E.Y., and Prakriya, M. (2012). Ion channels and transporters in lymphocyte function and immunity. Nature Reviews Immunology 2012 12:7 12, 532–547. 10.1038/nri3233.

88. Hogan, P.G., Lewis, R.S., and Rao, A. (2010). Molecular Basis of Calcium Signaling in Lymphocytes: STIM and ORAI. Annu. Rev. Immunol. 28, 491. 10.1146/ANNUREV.IMMUNOL.021908.132550.

89. Mazurek, M.P., Prasad, P.D., Gopal, E., Fraser, S.P., Bolt, L., Rizaner, N., Palmer, C.P., Foster, C.S., Palmieri, F., Ganapathy, V., et al. (2010). Molecular origin of plasma membrane citrate transporter in human prostate epithelial cells. EMBO Rep. 11, 431–437. 10.1038/EMBOR.2010.51/FIGURES/5.

90. Garcia, E., Connelly, M.A., Matyus, S.P., Otvos, J.D., and Shalaurova, I. (2021). High-throughput nuclear magnetic resonance measurement of citrate in serum and plasma in the clinical laboratory. Pract. Lab. Med. 25, e00213. 10.1016/J.PLABM.2021.E00213.

91. Costello, L.C., and Franklin, R.B. (2016). Plasma Citrate Homeostasis: How It Is Regulated; And Its Physiological and Clinical Implications. An Important, But Neglected, Relationship in Medicine. HSOA J. Hum. Endocrinol. 1, 005. 10.24966/he-9640/100005.

92. Dickens, F. (1941). The citric acid content of animal tissues, with reference to its occurrence in bone and tumour. Biochemical Journal 35, 1011. 10.1042/BJ0351011.

93. Kenny, A.D., Draskoczy, P.R., and Goldhaber, P. (1959). Citric acid production by resorbing bone in tissue culture. https://doi.org/10.1152/ajplegacy.1959.197.2.502 197, 502–504. 10.1152/AJPLEGACY.1959.197.2.502.

94. Schwarcz, H.P., Agur, K., and Jantz, L.M. (2010). A New Method for Determination of Postmortem Interval: Citrate Content of Bone*. J. Forensic Sci. 55, 1516–1522. 10.1111/J.1556-4029.2010.01511.X.

95. Knuuttila, M., Lappalainen, R., Alakuijala, P., and Lammi, S. (1985). Statistical evidence for the relation between citrate and carbonate in human cortical bone. Calcif. Tissue Int. 37, 363–366. 10.1007/BF02553702.

96. Taylor, T.G. (1960). The nature of bone citrate. Biochim. Biophys. Acta 39, 148–149. 10.1016/0006-3002(60)90131-1.

97. Hu, Y.Y., Rawal, A., and Schmidt-Rohr, K. (2010). Strongly bound citrate stabilizes the apatite nanocrystals in bone. Proc. Natl. Acad. Sci. U. S. A. 107, 22425–22429. 10.1073/PNAS.1009219107/-/DCSUPPLEMENTAL.

98. Simpson, D.P. (1983). Citrate excretion: a window on renal metabolism. Am. J. Physiol. 244. 10.1152/AJPRENAL.1983.244.3.F223.

99. Baruch, S.B., Burich, R.L., Eun, C.K., and King, V.F. (1975). Renal Metabolism of Citrate. Medical Clinics of North America 59, 569–582. 10.1016/S0025-7125(16)32009-0.

100. Costello, L.C., and Franklin, R.B. (1991). Concepts of citrate production and secretion by prostate 1. Metabolic relationships. Prostate 18, 25–46. 10.1002/PROS.2990180104.

101. Kline, E.E., Treat, E.G., Averna, T.A., Davis, M.S., Smith, A.Y., and Sillerud, L.O. (2006). Citrate Concentrations in Human Seminal Fluid and Expressed Prostatic Fluid Determined via 1H Nuclear Magnetic Resonance Spectroscopy Outperform Prostate Specific Antigen in Prostate Cancer Detection. J. Urol. 176, 2274–2279. 10.1016/J.JURO.2006.07.054.

102. Chen, J., Xu, Z., Zhao, H., and Jiang, X. (2007). Citrate in expressed prostatic secretions has the feasibility to be used as a useful indicator for the diagnosis of category IIIB prostatitis. Urol. Int. 78, 230–234. 10.1159/000099343.

103. Mann, T., and Lutwak-Mann, C. (1981). Male Reproductive Function and Semen. Male Reproductive Function and Semen. 10.1007/978-1-4471-1300-3.

104. Salisbury, G.W. (1965). Mammalian Reproduction: The Biochemistry of Semen and of the Male Reproductive Tract. Thaddeus Mann. Methuen, London; Wiley, New York, 1964. xxiv + 493 pp. Illus. $16.50. Science (1979). 147, 727–728. 10.1126/SCIENCE.147.3659.727.B.

105. Glusker, J.P. (1971). 14 Aconitase. Enzymes (Essen). 5, 413–439. 10.1016/S1874-6047(08)60097-9.

106. Krebs, H.A. (1953). The equilibrium constants of the fumarase and aconitase systems. Biochemical Journal 54, 78. 10.1042/BJ0540078.

107. Costello, L.C., Liu, Y., Franklin, R.B., and Kennedy, M.C. (1997). Zinc inhibition of mitochondrial aconitase and its importance in citrate metabolism of prostate epithelial cells. Journal of Biological Chemistry 272, 28875–28881. 10.1074/jbc.272.46.28875.

108. Costello, L.C., Franklin, R.B., Liu, Y., and Kennedy, M.C. (2000). Zinc causes a shift toward citrate at equilibrium of the m-aconitase reaction of prostate mitochondria. J. Inorg. Biochem. 78, 161–165. 10.1016/S0162-0134(99)00225-1.

109. Dirckx, N., Zhang, Q., Chu, E.Y., Tower, R.J., Li, Z., Guo, S., Yuan, S., Khare, P.A., Zhang, C., Verardo, A., et al. (2022). A specialized metabolic pathway partitions citrate in hydroxyapatite to impact mineralization of bones and teeth. Proc. Natl. Acad. Sci. U. S. A. 119, e2212178119. 10.1073/PNAS.2212178119/-/DCSUPPLEMENTAL.

110. Franklin, R.B., Chellaiah, M., Zou, J., Reynolds, M.A., and Costello, L.C. (2014). Evidence that Osteoblasts are Specialized Citrate-producing Cells that Provide the Citrate for Incorporation into the Structure of Bone. Open Bone J. 6, 1. 10.2174/1876525401406010001.

111. Franklin, R.B., Ma, J., Zou, J., Guan, Z., Kukoyi, B.I., Feng, P., and Costello, L.C. (2003). Human ZIP1 is a major zinc uptake transporter for the accumulation of zinc in prostate cells. J. Inorg. Biochem. 96, 435. 10.1016/S0162-0134(03)00249-6.

112. McCormack, J.G., and Denton, R.M. (1979). The effects of calcium ions and adenine nucleotides on the activity of pig heart 2-oxoglutarate dehydrogenase complex. Biochem. J. 180, 533–544. 10.1042/BJ1800533.

113. Denton, R.M. (2009). Regulation of mitochondrial dehydrogenases by calcium ions. Biochimica et Biophysica Acta (BBA) - Bioenergetics 1787, 1309–1316. 10.1016/J.BBABIO.2009.01.005.

114. Pettit, F.H., Roche, T.E., and Reed, L.J. (1972). Function of calcium ions in pyruvate dehydrogenase phosphatase activity. Biochem. Biophys. Res. Commun. 49, 563–571. 10.1016/0006-291X(72)90448-2.

115. Johnson, E.N., Lee, Y.M., Sander, T.L., Rabkin, E., Schoen, F.J., Kaushal, S., and Bischoff, J. (2003). NFATc1 mediates vascular endothelial growth factor-induced proliferation of human pulmonary valve endothelial cells. J. Biol. Chem. 278, 1686–1692. 10.1074/JBC.M210250200.

116. Jang, G.H., Park, I.S., Yang, J.H., Bischoff, J., and Lee, Y.M. (2010). Differential function of genes regulated by VEGF-NFATc1 signaling pathway in migration of pulmonary valve endothelial cells. FEBS Lett. 584, 141. 10.1016/J.FEBSLET.2009.11.031.

117. Suehiro, J.I., Kanki, Y., Makihara, C., Schadler, K., Miura, M., Manabe, Y., Aburatani, H., Kodama, T., and Minami, T. (2014). Genome-wide Approaches Reveal Functional Vascular Endothelial Growth Factor (VEGF)-inducible Nuclear Factor of Activated T Cells (NFAT) c1 Binding to Angiogenesis-related Genes in the Endothelium. J. Biol. Chem. 289, 29044. 10.1074/JBC.M114.555235.

118. Schwieger, M., Schüler, A., Forster, M., Engelmann, A., Arnold, M.A., Delwel, R., Valk, P.J., Löhler, J., Slany, R.K., Olson, E.N., et al. (2009). Homing and invasiveness of MLL/ENL leukemic cells is regulated by MEF2C. Blood 114, 2476–2488. 10.1182/BLOOD-2008-05-158196.

119. Homminga, I., Pieters, R., Langerak, A.W., de Rooi, J.J., Stubbs, A., Verstegen, M., Vuerhard, M., Buijs-Gladdines, J., Kooi, C., Klous, P., et al. (2011). Integrated transcript and genome analyses reveal NKX2-1 and MEF2C as potential oncogenes in T cell acute lymphoblastic leukemia. Cancer Cell 19, 484–497. 10.1016/J.CCR.2011.02.008.

120. Pon, J.R., and Marra, M.A. (2015). MEF2 transcription factors: developmental regulators and emerging cancer genes. Oncotarget 7, 2297. 10.18632/ONCOTARGET.6223.

121. Van Anken, E., Romijn, E.P., Maggioni, C., Mezghrani, A., Sitia, R., Braakman, I., and Heck, A.J.R. (2003). Sequential waves of functionally related proteins are expressed when B cells prepare for antibody secretion. Immunity 18, 243–253. 10.1016/S1074-7613(03)00024-4.

122. Chen, J.L., Ahluwalia, J.P., and Stamnes, M. (2002). Selective effects of calcium chelators on anterograde and retrograde protein transport in the cell. J. Biol. Chem. 277, 35682–35687. 10.1074/JBC.M204157200.

123. Kupfer, A., Dennert, G., and Singer, S.J. (1983). Polarization of the Golgi apparatus and the microtubule-organizing center within cloned natural killer cells bound to their targets. Proc. Natl. Acad. Sci. U. S. A. 80, 7224–7228. 10.1073/pnas.80.23.7224.

124. Bunnell, S.C., Kapoor, V., Trible, R.P., Zhang, W., and Samelson, L.E. (2001). Dynamic actin polymerization drives T cell receptor-induced spreading: a role for the signal transduction adaptor LAT. Immunity 14, 315–329. 10.1016/S1074-7613(01)00112-1.

125. Kwiatkowski, D.J. (1999). Functions of gelsolin: motility, signaling, apoptosis, cancer. Curr. Opin. Cell Biol. 11, 103–108. 10.1016/S0955-0674(99)80012-X.

126. Benitah, J.P., Pereira, L., Perrier, R., Mercadier, J.J., Sabourin, J., and Gómez, A.M. (2026). Local calcium dynamics and signalling in cardiomyocytes. Nature Reviews Cardiology 2026, 1–16. 10.1038/s41569-026-01286-8.

127. Ebashi, S. (1972). Calcium Ions and Muscle Contraction. Nature 1972 240:5378 240, 217–218. 10.1038/240217a0.

128. Groigno, L., and Whitaker, M. (1998). An anaphase calcium signal controls chromosome disjunction in early sea urchin embryos. Cell 92, 193–204. 10.1016/S0092-8674(00)80914-9.

129. Ciapa, B., Pesando, D., Wilding, M., and Whitaker, M. (1994). Cell-cycle calcium transients driven by cyclic changes in inositol trisphosphate levels. Nature 368, 875–878. 10.1038/368875A0.

130. Pinton, P., Giorgi, C., Siviero, R., Zecchini, E., and Rizzuto, R. (2008). Calcium and apoptosis: ER-mitochondria Ca2+ transfer in the control of apoptosis. Oncogene 2008 27:50 27, 6407–6418. 10.1038/onc.2008.308.

131. Orrenius, S., Zhivotovsky, B., and Nicotera, P. (2003). Regulation of cell death: the calcium–apoptosis link. Nature Reviews Molecular Cell Biology 2003 4:7 4, 552–565. 10.1038/nrm1150.

132. Lewis, C.A., Parker, S.J., Fiske, B.P., McCloskey, D., Gui, D.Y., Green, C.R., Vokes, N.I., Feist, A.M., Vander Heiden, M.G., and Metallo, C.M. (2014). Tracing compartmentalized NADPH metabolism in the cytosol and mitochondria of mammalian cells. Mol. Cell 55, 253–263. 10.1016/J.MOLCEL.2014.05.008.

133. Lien, E.C., Vu, N., Westermark, A.M., Danai, L. V., Lau, A.N., Gültekin, Y., Kukurugya, M.A., Bennett, B.D., and Vander Heiden, M.G. (2023). Effects of aging on glucose and lipid metabolism in mice. bioRxiv, 2023.12.17.572088. 10.1101/2023.12.17.572088.

134. Love, M.I., Huber, W., and Anders, S. (2014). Moderated estimation of fold change and dispersion for RNA-seq data with DESeq2. Genome Biol. 15. 10.1186/S13059-014-0550-8.

135. Türei, D., Schaul, J., Palacio-Escat, N., Bohár, B., Bai, Y., Ceccarelli, F., Çevrim, E., Daley, M., Darcan, M., Dimitrov, D., et al. (2026). OmniPath: integrated knowledgebase for multi-omics analysis. Nucleic Acids Res. 54, D652–D660. 10.1093/NAR/GKAF1126.

136. Ge, S.X., Jung, D., Jung, D., and Yao, R. (2020). ShinyGO: a graphical gene-set enrichment tool for animals and plants. Bioinformatics 36, 2628–2629. 10.1093/BIOINFORMATICS/BTZ931.

137. Agrawal, S., Kumar, S., Sehgal, R., George, S., Gupta, R., Poddar, S., Jha, A., and Pathak, S. (2019). El-MAVEN: A Fast, Robust, and User-Friendly Mass Spectrometry Data Processing Engine for Metabolomics. Methods Mol. Biol. 1978, 301–321. 10.1007/978-1-4939-9236-2_19.

138. Cerami, E., Gao, J., Dogrusoz, U., Gross, B.E., Sumer, S.O., Aksoy, B.A., Jacobsen, A., Byrne, C.J., Heuer, M.L., Larsson, E., et al. (2012). The cBio cancer genomics portal: an open platform for exploring multidimensional cancer genomics data. Cancer Discov. 2, 401–404. 10.1158/2159-8290.CD-12-0095.

139. Honcharuk, V., Zainab, A., Horimoto, Y., Takemoto, K., Diez, D., Kawaoka, S., and Vandenbon, A. (2026). DeepSpaceDB: a spatial transcriptomics atlas for interactive in-depth analysis of tissues and tissue microenvironments. Nucleic Acids Res. 54, D1017–D1030. 10.1093/NAR/GKAF1117.

140. Hao, Y., Stuart, T., Kowalski, M.H., Choudhary, S., Hoffman, P., Hartman, A., Srivastava, A., Molla, G., Madad, S., Fernandez-Granda, C., et al. (2024). Dictionary learning for integrative, multimodal and scalable single-cell analysis. Nat. Biotechnol. 42, 293–304. 10.1038/S41587-023-01767-Y.

